# Bacteria deliver a microtubule-binding protein into mammalian cells to promote colonization

**DOI:** 10.1101/2025.06.17.660209

**Authors:** Michael S. Costello, Bryan C. Neumann, Bonnie J. Cuthbert, Jana Holubová, Mia W. Raimondi, Fernando Garza-Sánchez, Abdul Samad, Ladislav Bumba, Jacob A. Torres, Nickolas Holznecht, Jessica Mendoza, Ondrej Stanek, Sasiprapa Prombhul, Thomas Weimbs, Meghan A. Morrissey, Diego Acosta-Alvear, David A. Low, Peter Šebo, Celia W. Goulding, Shane Gonen, Christopher S. Hayes

## Abstract

Pathogenic *Bordetella* bacteria infect the ciliated respiratory epithelia of mammalian and avian hosts. Several bacterial proteins mediate host cell adhesion, but filamentous hemagglutinin (FhaB) is a principal adhesin because mutants lacking this protein exhibit profound colonization defects. Here, we show that FhaB carries a C-terminal microtubule-binding domain (FhaB-CT), which is translocated into the host-cell cytoplasm to promote bacterial colonization. Cryogenic electron microscopy of microtubule-bound FhaB-CT shows that the domain binds primarily to α-tubulin through a network of polar interactions. Live-cell microscopy of infected tracheal explants reveals that FhaB-CT delivery is required for *Bordetella* to occupy a niche at the base of cilia on airway epithelia. Finally, we demonstrate that the microtubule-binding domain is required for long-term colonization of the mouse nasal cavity by *B. pertussis*. These observations suggest that the FhaB-CT domain is delivered into motile cilia, where it interacts with axonemal microtubules. We propose that *Bordetella* initially adhere to the tips of cilia, then deploy multiple FhaB adhesin molecules to migrate to the base of the cilial forest. This mechanism enables *Bordetella* to resist removal by the mucociliary ‘escalator’ that clears the respiratory tract of microbes and debris.

---

Recent years have witnessed a resurgence in whooping cough, or pertussis, which is caused by *Bordetella pertussis* (*1*). These pathogenic bacteria colonize ciliated epithelia of the upper respiratory tract and elicit a highly contagious illness that often leads to serious complications. Following the COVID-19 pandemic, pertussis outbreaks reminiscent of the pre-vaccine era swept through the EU and other high-income countries (*2, 3*). Despite high immunization rates, pertussis remains the least-controlled vaccine-preventable infectious disease, with an estimated 24 million whooping cough cases and ∼160,000 pertussis-related infant deaths occurring annually worldwide (*4*). The increased incidence of pertussis and related morbidity are largely due to the failure of the acellular vaccine to confer long-term protection against infection and restrict pathogen transmission (*5, 6*). Current vaccines contain inactivated pertussis toxin and one or more cell-surface antigens including filamentous hemagglutinin (FHA, **Fig. 1A**) (*6*), an adhesin required for airway colonization (*7-9*). FHA is a processed form of the FhaB protein that lacks the C-terminal ‘prodomain’ region (**Fig. 1B**) (*10, 11*). The FhaB prodomain is subject to proteolytic degradation in the bacterial periplasm as the adhesin is exported by the outer-membrane localized FhaC transport protein (*12, 13*). FHA is then cleaved from the bacterial cell surface by other proteases (*14, 15*), and the released protein has been hypothesized to act as an extracellular matrix to promote host-cell adhesion and biofilm formation (*16-19*). Although processed FHA is widely considered to be the mature adhesin, here we demonstrate that full-length FhaB plays an essential role in colonization by translocating a unique microtubule-binding domain into host cells.

**Fig. 1.**
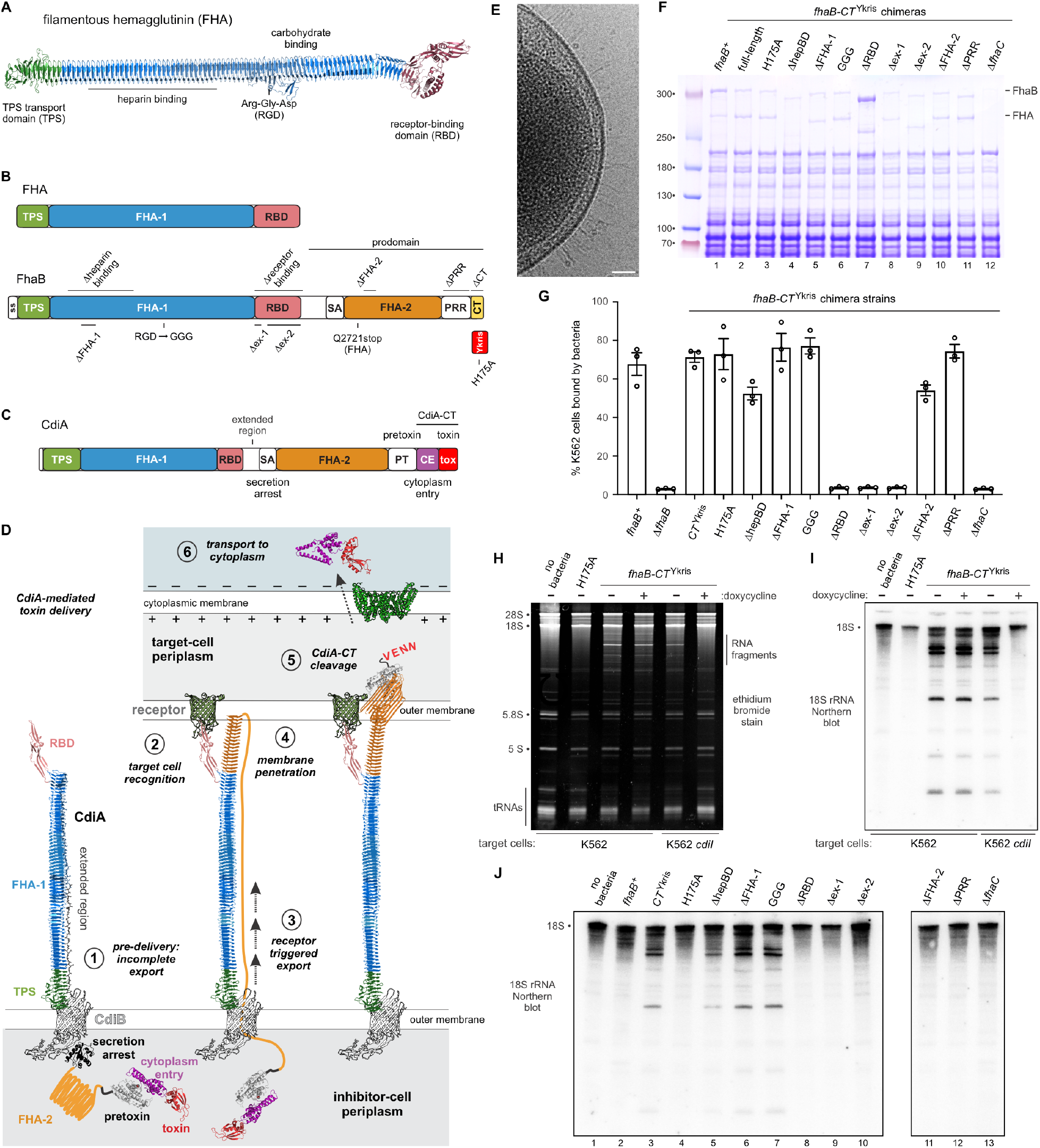
FhaB delivers its C-terminal domain into mammalian cells. (**A**) AlphaFold3 model of processed FHA. (**B**) Domain architectures of FHA and the FhaB precursor protein. (**C**) CdiA domain architecture. (**D**) Model of CdiA-mediated toxin delivery. (**E**) Electron cryotomography of cell-surface FhaB. (**F**) SDS-PAGE analysis of chimeric FhaB-CT^Ykris^ proteins. (**G**) Bacterial adhesion to K562 cells. The percentage of K562 bound to fluorescent bacteria was quantified by flow cytometric counting of 10,000 events. Reported values are averages ± SEM for three independent experiments. (**H**) Urea-PAGE analysis of RNA from K562 infections. (**I**) Northern blot analysis of 18S rRNA cleavage. Samples from panel h were electroblotted and hybridized with radiolabeled probes to human 18S rRNA. (**J**) Northern blot analysis of 18S rRNA cleavage mediated by FhaB-CT^Ykris^ chimeras.

## FhaB shares domain architecture with antibacterial CdiA effectors

FhaB and its FhaC transporter share several common features with the CdiA and CdiB proteins that mediate contact-dependent growth inhibition (CDI) (*20*). CDI is a mechanism of inter-cellular competition whereby Gram-negative bacteria deliver protein toxins directly into neighboring target cells (*21, 22*). CDI^+^ bacteria use CdiB transporters to present toxic CdiA effectors on the bacterial cell surface in the same manner that FhaC exports FhaB (**Fig. 1D**) (*23, 24*). Moreover, CdiA and FhaB have very similar domain architectures (**Figs. 1B & 1C**). Both proteins contain prominent FHA-1 peptide-repeat regions, which are predicted to fold into elongated β-helices (**Figs. 1A & 1D**) (*11, 25*). We previously showed that the FHA-1 domain of CdiA forms an extracellular filament that projects ∼33 nm from the bacterial surface (*26*), and electron cryotomography of *Bordetella bronchiseptica* reveals that FhaB is also presented as filaments on the cell surface (**Figs. 1E & S1**). The distal tip of the CdiA filament carries a receptor-binding domain (RBD) that mediates adhesion to target bacteria (**Figs. 1C & 1D**, step 2). Upon binding receptor, the C-terminal domains of CdiA are exported to deliver toxin into the target bacterium (**Fig. 1D**, step 3). The FHA-2 domain penetrates the target-cell outer membrane (**Fig. 1D**, step 4) and deposits the C-terminal toxin (CdiA-CT) into the periplasm through autoproteolysis (**Fig. 1D**, step 5) (*26-28*). The released toxin then hijacks receptors in the cytoplasmic membrane to enter the target-cell cytosol, where it gains access to its substrates (**Fig. 1D**, step 6) (*29*). AlphaFold3 (*30*) modeling indicates that CdiA and FhaB contain similar RBD structures built of two excursions from the β-helix (**Figs. 1A & S2**). Importantly, FhaB is also predicted to contain a domain with structural homology to FHA-2 of CdiA (**Fig. S3**). The C-terminus of FhaB diverges from CdiA and is composed of an unstructured prolyl-rich region (PRR) linked to a small C-terminal domain (FhaB-CT) of unknown function (**Figs. 1B & S4**).

Based on the overall structural similarity, we reasoned that FhaB functions like CdiA to translocate its C-terminal domain into host cells during colonization.

## FhaB translocates heterologous cargo into mammalian cells

The FhaB-CT domain has no known biochemical activity, so we first explored the CDI delivery hypothesis by testing whether FhaB can transfer a heterologous reporter into mammalian cells. We replaced the native FhaB-CT of *B. bronchiseptica* RB50 with the C-terminal RNase domain of CdiA from *Yersinia kristensenii* (CT^Ykris^) (**Fig. 1B**), because this nuclease is active after delivery into target bacteria (*31*). The FhaB-CT^Ykris^ fusion accumulates to somewhat lower levels than wild-type FhaB and is processed into FHA (**Fig. 1F**, lanes 1 & 2), suggesting that the chimeric protein is presented on the bacterial surface normally. To assess hemagglutination activity, we examined bacterial adhesion to the human K562 cell line (*32*), anticipating that these erythroleukemia cells express the receptor for FhaB. Indeed, bacteria expressing either wild-type FhaB or FhaB-CT^Ykris^ agglutinate K562 cells, whereas Δ*fhaB* mutants lacking the adhesin fail to bind (**Fig. S5**). Moreover, flow cytometry shows that FhaB and FhaB-CT^Ykris^ support the same level of bacterial adhesion to K562 cells (**Fig. 1G**), indicating that substitution of the native FhaB-CT with CT^Ykris^ does not interfere with hemagglutination activity.

We next tested whether the FhaB-CT^Ykris^ chimera delivers its RNase domain into the K562 cytoplasm. RNA isolated from K562 infected with *fhaB-CT*^Ykris^ bacteria contains several fragments that are not observed in uninfected cells (**Fig. 1H**). Northern blot analysis revealed that some of the fragments are derived from human 18S ribosomal RNA (**Fig. 1I**). These RNA species are presumably degradation products, because they do not accumulate when the RNase domain is inactivated with a His175Ala mutation in the catalytic site (**Figs. 1H & 1I**) (*31*). To demonstrate that the CT^Ykris^ domain is responsible for RNA degradation, we expressed the CdiI^Ykris^ immunity protein in K562 cells. CdiI immunity proteins are produced by all CDI^+^ bacteria to protect against auto-intoxication (*33*). As expected, nuclease activity is neutralized when K562 target cells express immunity protein from a doxycycline inducible promoter (**Figs. 1H & 1I**). Together, these data indicate that CT^Ykris^ is transferred into the K562 cytoplasm.

Although the CT^Ykris^ RNase domain enters the cytosol, delivery could be due to endocytosis or some other import pathway unrelated to the CDI mechanism. As predicted by the CDI hypothesis, we found that the FhaC transport protein is required for both bacterial adhesion (**Fig. 1G**) and RNase delivery (**Fig. 1J**, lane 13). However, because Δ*fhaC* mutants produce much lower levels of FhaB-CT^Ykris^ (**Figs. 1F & S6A**, lane 12), these results do not necessarily exclude alternative delivery pathways. Therefore, we deleted different regions of the FhaB-CT^Ykris^ chimera to interrogate adhesion and RNase delivery activities (**Figs. 1B & 1F**). FHA is known to bind a number of cell surface antigens including αβ integrins (*34*), heparan sulfate proteoglycans (*35*) and lactosyl ceramides (*36*), and the corresponding interaction sites map to the β-helical FHA-1 repeat region (**Fig. 1A**) (*37*). Deletion of a 98-residue segment from the β-helix (ΔFHA-1) has no effect on K562 adhesion (**Fig. 1G**) or RNase delivery (**Fig. 1J**, lane 6). More extensive deletion of the entire heparin-binding domain (ΔhepBD) diminishes, but does not abrogate, adhesion and RNase delivery (**Figs. 1G & 1J**, lane 5). Substitution of the integrin-binding Arg-Gly-Asp motif with Gly-Gly-Gly (GGG) has no effect on cell adhesion or RNase delivery (**Figs. 1G & 1J**, lane 7). These findings indicate that heparin and αβ integrins may stabilize FhaB-mediated adhesion, but they are not the surface receptors required for CT^Ykris^ delivery. By contrast, mutations that excise the predicted RBD (ΔRBD) or its constituent subdomains (Δex-1 and Δex-2) completely block adhesion and RNase delivery activities (**Figs. 1G & 1J**, lanes 8, 9 & 10). These data are consistent with the CDI mechanism, in which receptor recognition triggers CT translocation into the target cell (*26*). The CDI hypothesis also predicts that the membrane-penetrating FHA-2 domain should be required for RNase delivery, but not for adhesion to target cells (*26*). Accordingly, disruption of FHA-2 has modest effects on K562 adhesion, but completely abrogates RNase delivery (**Figs. 1G & 1J**, lane 11). Similarly, removal of the prolyl-rich region (ΔPRR) has no effect on bacterial adhesion, though this segment is essential for RNase translocation into K562 cells (**Figs. 1G & 1J**, lane 12). These results demonstrate that cell adhesion is necessary but not sufficient for CT^Ykris^ delivery. We repeated the adhesion and RNase delivery experiments with Vero cells to test whether our findings are applicable to other target cell lines. In general, we obtained similar results, though bacteria expressing the ΔhepBD variant exhibit decreased adhesion and fail to deliver RNase into Vero cells (**Figs. S6B, S6C & S6D**, lane 5), suggesting that heparan sulfate proteoglycans are important for adhesion to these particular cells. Overall, our findings indicate that FhaB uses a cell contact-dependent mechanism to translocate its C-terminus into the eukaryotic cytosol.

## Bordetella FhaB-CT binds to microtubules

Given that FhaB transfers a heterologous cargo into mammalian cells, we reasoned that its native C-terminus interacts with cytosolic factors in the host. To identify interacting partners, we fused GFP to the PRR-CT region of FhaB and expressed the fusion protein in U2OS cells. Confocal immunofluorescence microscopy revealed that GFP-PRR-CT forms cytoplasmic fibrils that co-localize with the microtubule network (**Fig. 2A & Movie S1**). This molecular association is direct because purified PRR-CT co-sediments with *in vitro* assembled microtubules during ultracentrifugation (**Fig. 2B**, lane 4). By contrast, the C-terminal regions of unrelated FhaB proteins from *Enterobacter* SST-3 and *Yersinia frederiksenii* ATCC 33641 do not bind microtubules and remain in the supernatant fraction in this assay (**Fig. 2B**, lanes 7 & 11). We speculated that the PRR may contribute to binding because Tau protein contains a similar prolyl-rich segment that influences microtubule interactions (*38*). Although truncation of the PRR appears to reduce binding affinity, the minimal FhaB-CT domain (residues Pro3610 - Lys3710) still interacts with microtubules *in vitro* (**Fig. 2C**, lane 12). Accordingly, GFP-CT fusions lacking the PRR also decorate the microtubule network when expressed in U2OS cells (**Movies S2 & S3**). Lastly, we confirmed direct interactions using negative-stain electron microscopy to visualize purified FhaB-CT domains bound to *in vitro* assembled microtubules (**Fig. S7**).

**Figure 2.**
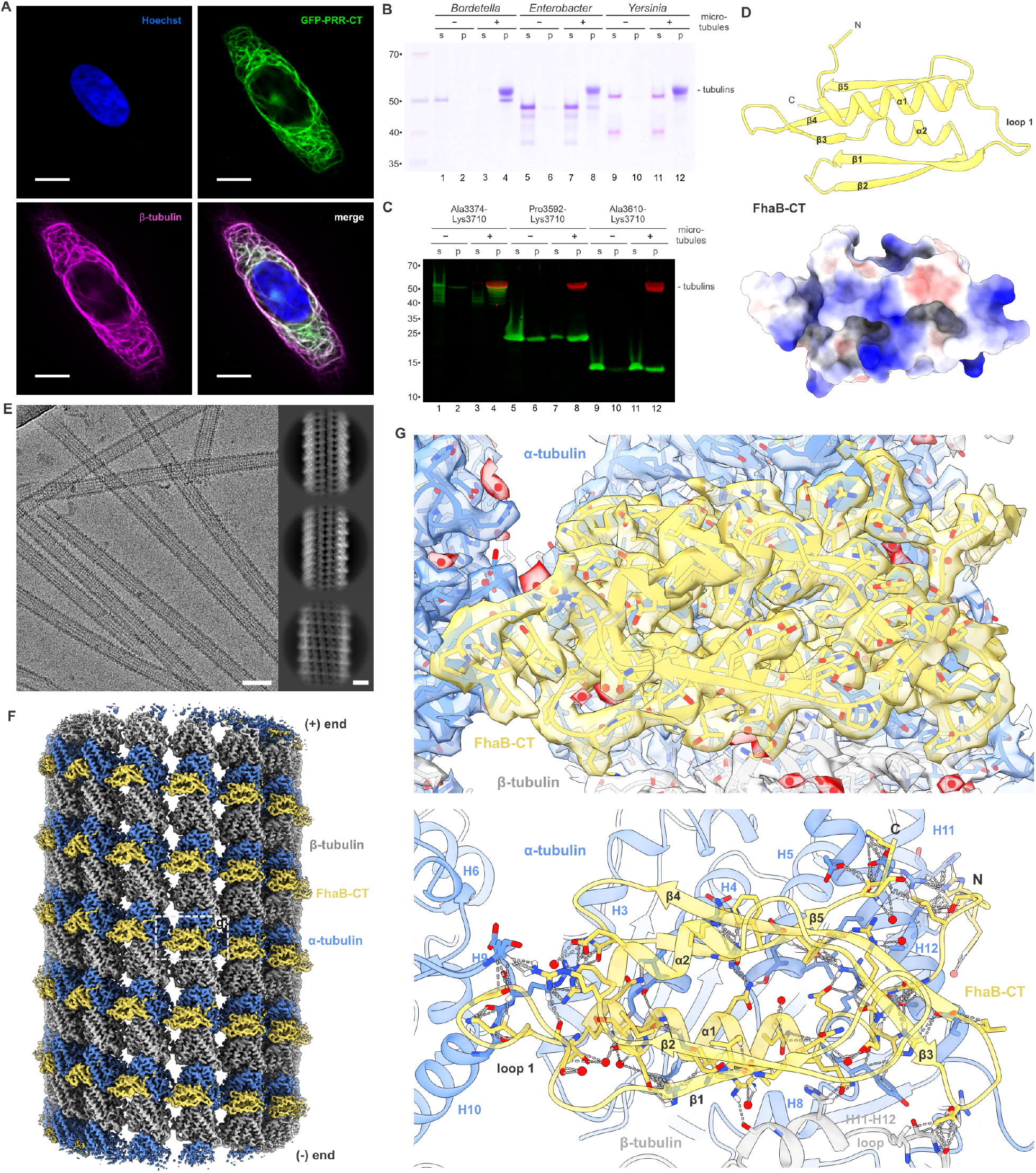
FhaB-CT binds to microtubules. (**A**) Ectopic expression of GFP-PRR-CT in U2OS cells. Transfected cells were stained with anti-β-tubulin and Hoechst to visualize microtubules and cell nuclei, respectively. Scale bar = 10 µm. (**b**) *In vitro* microtubule-binding assay. Purified FhaB fragments were centrifuged at 100,000 ×*g* with or without microtubules. Supernatant (s) and pellet (p) fractions were analyzed by SDS-PAGE. (**C**) FhaB-CT binds to microtubules. Purified His_6_-FhaB-CT constructs were centrifuged at 100,000 ×*g* with or without microtubules. Supernatant (s) and pellet (p) fractions were analyzed by SDS-PAGE and immunoblotting with anti-FhaB-CT antibodies. (**D**) Crystal structure of FhaB-CT. Negative and positive electrostatic surface potentials are indicated in red and blue, respectively. (**E**) Representative cryo-EM image of microtubules decorated with FhaB-CT domains (bar = 50 nm). Inset shows 2D classes (bars = 10 nm). (**F**) Full cryo-EM map of FhaB-CT domains bound to a 14-3 microtubule. (**G**) Refined model fit into the cryo-EM map (upper panel). Hydrogen-bonding network at the interface between FhaB-CT and the microtubule (lower panel).

To gain insight into this new microtubule-binding domain, we solved the crystal structure of FhaB-CT to 1.65 Å resolution (**Table S1**). FhaB-CT adopts an α/β fold that does not closely resemble any structures in the protein data bank, though its surface is electropositive as expected for a microtubule-binding protein (**Fig. 2D**). We used cryogenic electron microscopy to reconstruct a ∼2.4 Å resolution map of FhaB-CT bound to microtubules (**Figs. 2E, 2F, S8 & Table S2**). This high-resolution map enabled us to model FhaB-CT, α-tubulin and β-tubulin together with bound GTP/GDP nucleotides and taxol (**Figs. 2G & S9**). FhaB-CT domains decorate the outer surface of the microtubule (**Fig. 2F & Movie S4**), interacting primarily with α-tubulin through an extensive network of ion-pairs and hydrogen bonds (**Fig. 2G & Table S3**). Microtubule-bound FhaB-CT adopts essentially the same conformation as the crystal form (root mean square deviation 0.5 Å), though the β3-β4 hairpin is altered slightly likely due to interactions with β-tubulin. Loop 1 of FhaB-CT extends to interact with α-tubulin in the adjacent protofilament (**Figs. 2F, 2G & Movie S4**). In the 2.4 Å reconstruction, FhaB-CT appears to be absent from the microtubule seam, where α-tubulins interact with β-tubulins on the adjacent protofilament (**Movie S4**). For closer inspection, we aligned seam protofilaments from a subset of particles and were able to detect bound FhaB-CT in the newly aligned map (**Figs. S10, S11 & Movie S5**). However, the seam-bound domains are at lower resolution than FhaB-CT domains bound to other protofilaments. Together, these observations suggest that loop 1-mediated interactions bias FhaB-CT binding to non-seam protofilaments.

## The FhaB microtubule-binding domain promotes colonization at the base of cilia

The C-terminal microtubule-binding domain of FhaB is not strictly required for adhesion because chimeric FhaB-CT^Ykris^ supports bacterial binding to K562 and Vero cells. However, *Bordetella* normally colonize respiratory epithelia, which carry hundreds of motile cilia that beat in a concerted fashion to sweep microbes and debris from the airways (*39*). Because cilia movement is powered by the sliding motion of axonemal microtubules, these observations suggest that FhaB-CT is delivered to reinforce adhesion through interactions with the core axoneme. To explore this model, we used live-cell microscopy to monitor bacterial adhesion to respiratory epithelia in rat trachea explants. As expected, fluorescently labeled wild-type *fhaB*^*+*^ bacteria adhere avidly to ciliated epithelial cells (**Fig. 3A**). However, we found that Δ*fhaB* bacteria also bind cilia (**Fig. 3A**), which is somewhat surprising given that these mutants do not adhere to K562, Vero and A549 cell lines (**Figs. S12 & S13**). Presumably, Δ*fhaB* mutants bind using other adhesins, such as fimbriae, that are known to be produced during the *Bordetella* virulent growth phase (*40*).

**Figure 3.**
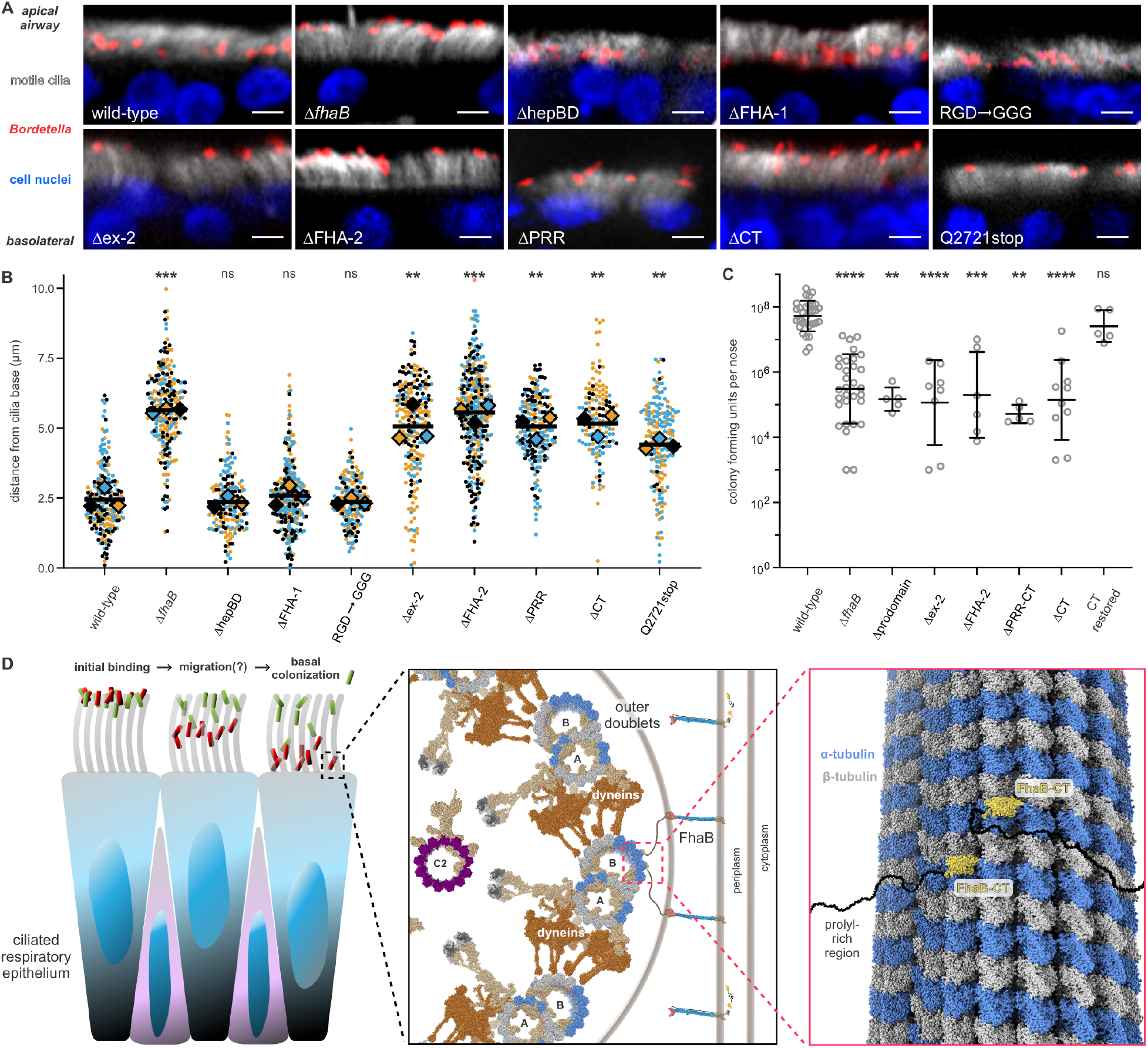
FhaB-CT delivery is required for colonization of host respiratory epithelia. (**A**) Live-cell microscopy of tdTomato-labeled RB50 bacteria bound to rat trachea epithelia. Cilia were stained with fluorescein-labeled taxol and nuclei visualized with Hoechst. Scale bars = 10 µm. (**B**) Live-cell microscopy of tdTomato-labeled RB50 bacteria bound to rat trachea epithelia. Cilia were stained with fluorescein-labeled taxol and nuclei visualized with Hoechst. Bars = 10 µm. (**C**) Quantification of bacterial distance to the base of cilia. Bacterial centroids were identified and to calculate the least Euclidean distance to the base of the cilial forest. Raw data and means (diamonds) are reported for three independent experiments using tracheas from different animals. Statistical significance was tested by parametric unpaired *t-*tests: *** (*p <* 0.001), ** (*p <* 0.01) and ns (not significant) (**D**) *B. pertussis* colonization of MyD88 mice. Nares of immunocompromised C57BL/6J MyD88 knockout mice were inoculated with 10^7^ cfu of *B. pertussis*, and cfu per nose were enumerated 7 d. Bars indicate the geometric mean ±?standard deviation. Statistical significance was assessed by one-way ANOVA followed by Dunnett’s multiple comparison test: **** (*p <* 0.0001), *** (*p <* 0.001), ** (*p <* 0.01) and ns (not significant). (**e**) Model for FhaB function during host colonization.

Although Δ*fhaB* mutants adhere to respiratory epithelia, their distribution in the cilia forest differs from wild-type *fhaB*^*+*^ bacteria. We quantified the relative positions of bound bacteria and found that wild-type *Bordetella* cluster near the base of cilia, whereas Δ*fhaB* mutant bacteria are found closer to the cilia tips (**Figs. 3A & 3B**). These results suggest that FhaB promotes bacterial penetration into the cilial forest. We next examined in-frame *fhaB* deletion strains to determine whether FhaB-CT delivery is required to occupy this niche. Deletions within the β-helical filament (ΔFHA-1 and ΔhepBD) have no effect on bacteria distribution (**Figs. 3A & 3B**). Similarly, mutants lacking the integrin-binding motif colonize like wild-type bacteria (**Figs. 3A & 3B**). However, deletion of excursion-2 from the RBD causes a colonization defect (**Figs. 3A & 3B**). Strains lacking the membrane penetrating FHA-2 domain and the PRR also phenocopy the cilia distribution of Δ*fhaB* null mutants (**Figs. 3A & 3B**), though the former mutants adhere to K562, Vero and A549 cells like wild-type bacteria (**Figs. S12 & S13**). These results strongly suggest that the microtubule-binding domain is required for normal colonization. Indeed, *fhaB(*Δ*CT)* bacteria lacking the microtubule-binding domain are unable to penetrate the cilia forest (**Figs. 3A & 3B**), but still adhere to cell lines like wild-type bacteria (**Figs. S12B & S13**). To assess whether prodomain processing is important for colonization, we introduced a stop codon at Gln2721 to mimic the processed FHA adhesin (**Figs. 1C**). This truncated FhaB is stable (**Fig. S12A**, lane 3) and is presented on the bacterial surface because it supports adhesion to K562, Vero and A549 cell lines (**Figs. S12b & S13**). However, bacteria lacking the FhaB prodomain do not colonize the base of cilia (**Figs. 3A & 3B**). Taken together, these results indicate that *Bordetella* relies on translocation of the C-terminal microtubule-binding domain to occupy its niche on ciliated respiratory epithelia.

Finally, we investigated the role of the microtubule-binding domain in host colonization using the human pathogen *B. pertussis*. FhaB proteins of *B. pertussis* and *B. bronchiseptica* are closely related and carry identical microtubule-binding domains (**Fig. S4**). Moreover, both *Bordetella* species rely on FhaB to colonize the murine respiratory tract (*7-9*). We recently showed that *B. pertussis* proliferates in the nasal cavities of homozygous MyD88 knock-out mice, which are defective for innate immune signaling through all but TLR3 Toll-like receptors (*9*). Following infection with 10^7^ cfu of *B. pertussis*, bacteria counts in the nasal cavity increase by an order of magnitude after 7 d, whereas Δ*fhaB* mutant populations decline ∼50-fold over the same time course (**Fig. 3C**). Strains producing in-frame deletion variants of FhaB show similar colonization defects, and strikingly the *fhaB(*Δ*CT)* mutants phenocopy the Δ*fhaB* null strain in this infection model (**Fig. 3C**). To rule out the possibility that the *fhaB(*Δ*CT)* strain acquired additional unintended mutations that adversely affect fitness in the host, we restored the FhaB-CT coding sequence and found that the complemented strain colonizes MyD88 knock-out mice like wild-type bacteria (**Fig. 3C**). These data demonstrate that delivery of the microtubule-binding FhaB-CT domain is critical for host colonization.

## Discussion

The mammalian respiratory tract is particularly challenging to colonize because its epithelium elaborates motile cilia that clear the airways of microbes and other debris (*39*). In addition, epithelial goblet cells produce a protective layer of mucous that ensnares bacteria for removal. Respiratory pathogens have long been known to deploy adhesins to resist elimination by this mucociliary ‘escalator’, but our results indicate that cell-surface interactions alone are not sufficient for *Bordetella* colonization. To overcome these physical obstacles, *Bordetella* species have repurposed a primordial inter-bacterial competition system that utilizes two distinct adhesive domains. FhaB interacts with cell-surface receptors like a conventional adhesin, but it also translocates its C-terminal microtubule-binding domain into cilia where it can bind the axoneme. In principle, the FhaB-CT domain could reinforce adhesion by anchoring bacteria to the internal microtubule core. However, we find that the microtubule-binding domain promotes bacterial colonization at the base of the cilial forest, suggesting that FhaB actively directs *Bordetella* to this location. This activity likely explains this adhesin’s importance for host colonization, because bacteria in this basal niche should be sheltered from the mucociliary escalator. Indeed, time-lapse microscopy indicates that bacteria at the tips of beating cilia are subject to greater displacement forces than those at the base (**Movies S6 & S7**). Moreover, in some instances, we observe bacteria being shed from cilia tips (**Movie S8**). Together, these findings show that FhaB is a critical colonization factor that enables *Bordetella* to penetrate the cilial forest to a protected niche.

How FhaB controls niche colonization remains to be determined. Presumably, *Bordetella* bacteria adhere initially to the cilia tips, then migrate down the organelles to the apical surface of the epithelium (**Fig. 3D**). According to this model, FhaB-mediated interactions must be reversible to allow bacteria to move to different positions along the cilium. This process could entail iterative rounds of adhesin attachment and release, with old binding interactions being relinquished as new contacts are established. Release of bound FhaB could be mediated by SphB1, which is a bacterial protease known to cleave FHA from the surface of *B. pertussis* (*14*). Bacterial migration would also require that FhaB-CT binding sites be readily available along the length of cilia. The motile cilium axoneme is composed of nine peripheral microtubule doublets that surround a central pair (**Fig. 3D**) (*41, 42*). Delivered FhaB-CT domains should be able to interact with peripheral doublets, because the flexible prolyl-rich region is long enough to span the ∼20-40 nm distance separating microtubules from the plasma membrane (**Fig. 3D**). FhaB-CT most likely interacts with B-tubules in the peripheral doublets, because the A-tubule surface is largely occluded by the outer-arm dynein complexes that power cilia movement (**Fig. 3D**). The migration model predicts that these dynein motors should not interfere with the binding of FhaB-CT to the microtubule doublets. Structural superimposition of dynein and FhaB microtubule-binding domains suggests that both proteins should be able to interact simultaneously (**Fig. S14**). Accordingly, FhaB does not appear to interfere with dynein function, because cilia continue to beat after *Bordetella* reach their basal niche (**Movie S9**). Another unresolved question concerns the driving force and directionality of bacterial migration. One intriguing possibility is that cilial motility energizes this process. Cilia beat at ∼10-20 Hz in a coordinated whip-like manner driven by dynein-mediated sliding of peripheral doublets past one another (*39*). It is possible that this sliding motion biases movement toward the cilium base. Alternatively, bacteria could ‘ping-pong’ between neighboring cilia during each recovery stroke to move incrementally closer to the base. Although the detailed mechanism remains obscure, FhaB clearly exploits the internal microtubule network to control bacterial position on the host cell surface.

## Acknowledgments

We thank the Advanced Light Source at Berkeley National Laboratories (ALS) and the Stanford Synchrotron Radiation Lightsource (SSRL) for help in data collection, and Rodger de Miranda for technical support.

## Funding

National Institutes of Health grant R01 GM117930 (CSH)

National Institutes of Health grant R21 AI151728 (DAL, DAA)

National Institutes of Health grant R35 GM142797 (SG)

National Institutes of Health grant U24 AG079683 (SG)

National Institutes of Health grant R21 AI185695 (CWG, CSH, SG)

Czech Science Foundation GA22-23578S (LB)

Czech Science Foundation 25-18104X (PS)

## Author contributions

Conceptualization: MSC, CSH

Methodology: MSC, BCN, BJC, JH, FGS, AS, LB, JAT, NH, JM, SP, MAM, PS, CSH

Investigation: MSC, BCN, BJC, JH, MWR, FGS, AS, LB, OS, CSH

Visualization: MSC, BCN, JH, MWR, FGS, AS, LB, CWG, SG, CSH

Funding acquisition: LB, DAA, DAL, PS, CWG, SG, CSH

Project administration: PS, CWG, SG, CSH

Supervision: PS, CWG, SG, CSH Writing – original draft: MSC, CSH

Writing – review & editing: MSC, BCN, BJC, FGS, TW, MAM, DAA, DAL, PS, CWG, SG, CSH

## Competing interests

The authors declare that they have no competing interests.

## Data and materials availability

The cryoEM maps from this study have been deposited to the Electron Microscopy Data Bank (EMDB) under accession codes EMD-49577 [https://www.ebi.ac.uk/pdbe/entry/emdb/EMD-49577] (*Bordetella* filamentous hemagglutinin (FhaB) C-terminal domain bound to microtubules), EMD-49766 [https://www.ebi.ac.uk/pdbe/entry/emdb/EMD-49766] (*Bordetella* filamentous hemagglutinin (FhaB) C-terminal domain bound to microtubules, ab initio), EMD-49767 [https://www.ebi.ac.uk/pdbe/entry/emdb/EMD-49767] (*Bordetella* filamentous hemagglutinin (FhaB) C-terminal domain bound to 14-3 microtubules), EMD-49768 [https://www.ebi.ac.uk/pdbe/entry/emdb/EMD-49768] (*Bordetella* filamentous hemagglutinin (FhaB) C-terminal domain bound to 14-3 microtubules, aligned), and EMD-49769 [https://www.ebi.ac.uk/pdbe/entry/emdb/EMD-49769] (*Bordetella* filamentous hemagglutinin (FhaB) C-terminal domain bound to microtubules at the seam). Atomic coordinates associated with this study have been deposited to the Protein Data Bank (PDB) under the accession codes 8SX0 [https://doi.org/10.2210/pdb8SX0/pdb] (*Bordetella* filamentous hemagglutinin (FhaB) C-terminal domain) and 9NNL [https://doi.org/10.2210/pdb9NNL/pdb] (*Bordetella* filamentous hemagglutinin (FhaB) C-terminal domain bound to microtubules). Previously reported cryoEM maps and models referred to in this manuscript can be found under accession codes EMD-5192, EMD-5195, EMD-8997, EMD-8998 and 7TQY. All other data are available in the main text or extended data.

## Notes

### Competing Interest Statement

The authors have declared no competing interest.

