## Supplemental Information for "Bacteria deliver a microtubule-binding protein into mammalian cells to promote colonization"

### Materials and Methods

**Bacterial strains and growth.** Bacterial strains are described in **Table S4**. *E. coli* bacteria were cultured in lysogeny broth (LB) or on LB agar. *B. bronchiseptica* RB50 was obtained from the American Type Culture Collection (ATCC), and *B. pertussis* Tohama I was obtained from the Culture Collection of the Institute Pasteur (catalogue No. CIP 81.32). *B. bronchiseptica* strains were cultured in either LB or Stainer-Scholte media or on Bordet-Gengou (BG) agar supplemented with 10% defibrinated sheep blood. *B. pertussis* strains were cultured in Stainer-Scholte medium or on BG agar supplemented with 1% glycerol and 15% defibrinated sheep blood. Where appropriate, media were supplemented with antibiotics at the following concentrations: ampicillin (Amp) 150 µg $mL^{-1}$, kanamycin (Kan) 50 µg $mL^{-1}$, streptomycin (Str) 50 µg $mL^{-1}$ and trimethoprim (Tp) 100 µg $mL^{-1}$. pSS4245 derived constructs were introduced into *Bordetella* by conjugation with either *E. coli* MFD (*43*) or *E. coli* SM10 λpir cells. *Bordetella* and *E. coli* bacteria were cocultured on BG agar supplemented with 10% defibrinated sheep blood and 50 mM $MgSO_4$ overnight at 37 °C. Transconjugants were selected on BG agar supplemented with 50 mM $MgSO_4$ and 50 µg/mL Kan. Allelic exchange was induced by plating onto BG agar without MgSO4. Surviving clones were screened by colony PCR to identify deletion strains. Plasmids pCH1252 (tdTomato) and pCH1509 (mNeon) were introduced into RB50 strains by conjugation with *E. coli* MFD cells, followed by selection on LB agar supplemented with Tp and 50 mM $MgSO_4$.

**Plasmid constructs.** Plasmids and oligonucleotides used in this study are listed in **Tables S5** & **S6**, respectively. RB50 deletion constructs were generated by PCR amplification of upstream and downstream homology fragments, which were ligated sequentially to plasmid pSS4245 (*44*). Plasmid pCH752 (Δ*fhaB*) was generated with products amplified by primers MSC66/MSC67 (NotI/SacI) and MSC68/MSC69 (SacI/BamHI). The *CT/cdiI*$^{\mathrm{Ykris}}$ module was amplified from plasmid pCH12241 (*31*) with primers CH5861/CH5862 (KpnI/SacI) and fused to an *fhaB* fragment amplified with primers CH5676/CH5677 (XbaI/KpnI). The resulting XbaI/SacI fragment was subcloned into SpeI/SacI digested pCH752 to generate plasmid pCH763 (*fhaB-CT/cdiI*$^{\mathrm{Ykris}}$). Plasmid pCH1326 (Δ*fhaC*) was generated with primers CH5807/CH5808 (SpeI/SacI) and CH5809/CH5810 (SacI/BamHI). Plasmid pCH1353 (Δ*fimABCD*) was generated with primers CH5502/CH5718 (SpeI/SacI) and CH5789/CH5790 (SacI/BamHI). Plasmid pCH1330 (Δ*fhaB-fimABCD-fhaC*) was generated with primers MSC66/MSC67 (NotI/SacI) and CH5809/CH5810 (SacI/BamHI). Plasmid pCH1536 (Δ*cyaA*) was generated with CH6121/CH6122 (SpeI/SacI) and CH6123/CH6124 (SacI/BamHI).

Plasmid pCH1320 (ΔFHA-1) was generated with primers CH5857/CH5858 (NotI/EcoRI) and CH5859/CH5869 (EcoRI/BamHI). Plasmid pCH1637 (ΔhepBD) was generated with primers CH5857/CH6244 (NotI/BamHI) and CH6245/CH6246 (BamHI/AvrII). Plasmid pCH1319 (ΔHis1147-Gly1377) was generated with CH5861/CH5862 (SpeI/EcoRI) and CH5863/CH5864 (EcoRI/SbfI). Plasmid pCH2479 (RGD→GGG) was generated with CH5861/CH5928 and CH5929/CH5864 and Gibson assembly into SpeI/SbfI digested pSS4245. Plasmid pCH1638

(Δex-1) was generated with CH6259/CH6260 (SpeI/SacI) and CH6261/CH6262 (SacI/EcoRI). Plasmid pCH1354 (Δex-2) was generated with CH5811/CH5812 (SpeI/SacI) and CH5813/CH5814 (SacI/EcoRI). Plasmid pCH1543 (ΔRBD) was generated with CH6259/CH6260 (SpeI/SacI) and CH5813/CH5814 (SacI/EcoRI). Plasmid pCH1170 (Gln2721stop) was generated with CH5775/CH5776 (XbaI/SacI) and MSC68/MSC69 (SacI/BamHI). Plasmid pCH1318 (ΔFHA-2) was generated with CH6214/CH6215 (NotI/SacI) and CH6216/CH6217 (SacI/BamHI). Plasmid pCH1352 (ΔPRR) was generated with CH5676/CH5720 (XbaI/KpnI) and CH5717/CH5718 (KpnI/SacI). Plasmid pCH2287 (ΔPRR-Ykris) was generated using primers CH5676/CH6325 and CH6328/CH5682. The fragments were combined using Gibson assembly, digested with XbaI/SacI and ligated to SpeI/SacI digested pSS4245. The downstream fragment was generated with primers MSC68/MSC69 (SacI/BamHI). Plasmid pCH1565 (ΔCT) was generated with CH6218/CH6219 (NotI/SacI) and MSC68/MSC69 (SacI/BamHI). The Δ*fhaB* deletion construct for *B. pertussis* Tohama I were generated by sequential ligation of products amplified with Bp-ΔfhaB-Not-for/Bp-ΔfhaB-Spe-rev and Bp-ΔfhaB-Spe-for/Bp-ΔfhaB-Bam-rev into pSS4245. All other Tohama I deletion constructs were generated by three-piece Gibson assembly of PCR products into SpeI/EcoRI digested plasmid pSS4245.

A fragment encoding the TPS transport domain of FhaB (residues Gly73 - Glu440) was amplified with CH5498/CH5499 and ligated to pET21d using NcoI/XhoI restriction sites to generate pCH1847. The 3′-coding region of *fhaB* was PCR amplified with primer pairs CH5502/CH5503, CH5671/CH5503 and CH5669/CH5503, and the products ligated to pSH21 via SpeI/XhoI restriction sites to generate FhaB-CT over-production plasmids pCH1811, pCH1165 and pCH1166, respectively. The FhaB-CT$^{SST3}$ coding region (corresponding to Ala2461 – Ser2752; NCBI RefSeq: WP_008499743.1) was amplified with CH5622/CH5623, digested with SpeI/XhoI and ligated to pSH21 to generate plasmid pCH1784. The FhaB-CT$^{Yfred}$ region (corresponding to Gly2409 – Arg2674; NCBI RefSeq: WP_050504536.1) was amplified with CH5624/CH5625, digested with SpeI/XhoI and ligated to pSH21 to generate plasmid pCH1785. The RB50 *ftsZAQ* operon was amplified with primers F-FtsZ/R-FtsZ and combined with a fragment containing the *E. coli ara* promoter (amplified with F-pBAD/R-pBAD) by Gibson assembly to generate plasmid pBBR1-*ftsZAQ*.

The $P_{trc}$ promoter region was amplified with CH5605/CH824 from pTrc99A and ligated to plasmid pSCBAD using NsiI/HindIII restriction sites to generate pBBR1. The coding sequence for tdTomato was amplified with MSC108/MSC109 and introduced into NcoI-XhoI digested pBBR1 by Gibson assembly to generate plasmid pCH1252. The coding sequence for mNeon was amplified with CH5936/CH5937 and CH5938/CH5939 and the products combined by Gibson assembly to remove the internal NcoI site. The assembled product was ligated to NcoI/XhoI digested pBBR1 to generate pCH1509.

The *cdiI*$^{Ykris}$ gene was amplified from pCH12241(*31*) with primers MSC81/MSC273 and ligated to KpnI/NotI digested pFBK001 to generate plasmid pMSC218. pMSC218 was used as a donor for the destination plasmid pINDUCER21 (*45*) (https://www.addgene.org/46948/) in a LR

Clonase™ reaction (Thermo Fisher Scientific) to produce plasmid pMSC224. For ectopic expression of GFP-FhaB-CT fusions in mammalian cells, FLAG-eGFP was amplified from DAA-A361 using oligonucleotides MSC37/MSC38, and *fhaB* fragments amplified with primers MSC299/MSC33, MSC39/MSC33 and MSC40/MSC33. Fragments were combined by Gibson assembly into AgeI/NotI-digested pFBK003 to generate entry plasmids. The constructs were used as donors in LR Clonase™ reactions with the pCW57.1 destination plasmid (https://www.addgene.org/41393/) to generate plasmids pMSC263, pMSC156 and pMSC158, respectively.

**Protein purification, antisera generation and immunoblotting.** N- and C-terminal domains of FhaB proteins were overproduced in *E. coli* strain CH2016 (*46*). Overnight cultures were diluted 1:10 into 15 mL of fresh LB media supplemented with 150 µg/ml ampicillin and incubated at 37 °C with shaking for 3 h. Protein expression was induced with 1 mM isopropyl-β-D-thiogalactopyranoside (IPTG) for 2 h. Cells were harvested by centrifugation and resuspended in 1 mL of urea lysis buffer [8 M urea, 20 mM Tris-HCl (pH 8.0), 150 mM NaCl] and subjected to one freeze-thaw cycle. Cell lysates were centrifuged at 20,000 ×*g* for 15 min at ambient temperature, and 100 µL of $Ni^{2+}$-nitrilotriacetic acid ($Ni^{2+}$-NTA) agarose resin was added to each clarified lysate. After incubation for 1 h, resins were collected by centrifugation and batch washed five times with 1.0 mL of urea lysis buffer supplemented with 20 mM imidazole. Proteins were eluted in urea lysis buffer supplemented with 50 mM EDTA. Purified FhaB-CT proteins (from plasmids pCH1784, pCH1785, pCH1811, pCH1865 and pCH1866) were dialyzed against 20 mM HEPES (pH 7.0), 20 mM NaCl for use in microtubule binding assays. The N-terminal TPS transport (pCH1847) and CT (pCH1811) domains from FhaB from were dialyzed against water and lyophilized for injection into rabbits to generate polyclonal antisera (Cocalico Biologicals Inc., Denver, PA, USA).

FhaB and FhaB-$CT^{Ykris}$ variants were analyzed by SDS-PAGE and immunoblotting. Overnight RB50 cultures were resuspended at an optical density at 600 nm ($OD_{600}$) of 0.4 in fresh LB media and incubated at 37 °C for 4 h. Bacteria were collected by centrifugation at 21,130 ×*g* in a microfuge and cell pellets frozen at -80 °C. Frozen pellets were resuspended in urea lysis buffer and broken by a second freeze-thaw cycle. Lysates were clarified by centrifugation at 21,130 ×*g* in a microfuge. Protein concentrations were estimated using Bradford reagent and equal amounts were analyzed by SDS-PAGE at 110 V using 7% polyacrylamide gels buffered with Tris-tricine. High-speed pellet and supernatant fractions from microtubule-binding assays were resolved by SDS-PAGE using 10% polyacrylamide gels buffered with Tris-tricine. For immunoblotting, gels were soaked for 10 min in 25 mM Tris, 192 mM glycine (pH 8.6), 10% methanol before transfer to polyvinylidene fluoride membranes at 17 V for 30 min using a semi-dry electroblotting apparatus. Membranes were blocked with 0.1% bovine serum albumin (BSA) in phosphate buffered saline (PBS) for 30 min at ambient temperature. Blots were incubated with rabbit anti-TPS transport domain, rabbit anti-FhaB-CT, or mouse anti-α-tubulin (Cell Signaling, 3873) antibodies at 1:5,000 dilutions in 0.1% BSA, PBS. After three 10 min washes with PBS, membranes were incubated with IRDye® 800CW-conjugated goat anti-rabbit IgG (LI-COR, 1:15,000 dilution) and/or IRDye® 680LT-conjugated goat anti-mouse IgG (LI-COR, 1:15,000) in PBS for 15 min. Immunoblots were visualized using a LI-COR Odyssey infrared imager.

**Bacterial adhesion.** K562 cells were seeded in antibiotic-free media at $2 \times 10^5$ cells/mL (K562) in 24-well plates. Overnight RB50 cultures were diluted 1:10 into fresh LB media and grown for 4 h at 37 °C to $OD_{600}$ ~ 2.5. Bacteria were diluted into antibiotic-free media and added to mammalian cells at a multiplicity of infection (MOI) of 100 bacteria per mammalian cell and gently swirled. Bacteria-cell mixtures were centrifuged at 500 ×*g* for 5 min, then incubated for 30 min at 37 °C with 5% $CO_2$. K562-bacteria suspensions were mixed gently, transferred to microfuge tubes and analyzed by flow cytometry. Cell binding was measured using a BD-Accuri C6 flow cytometer and CFlow plus software by analysis of 10,000 events after doublet discrimination. Uninfected K562 cells were used to establish an FL1-A gate.

Adherent cell lines were seeded at $1.5 \times 10^5$ cells/mL (Vero) or $7.5 \times 10^4$ cells/mL (A549) in 96 well glass-bottom plates and maintained in DMEM supplemented with 10% FBS. After 12 h, mNeon-labeled RB50 strains were added at a MOI of 100, centrifuged at 500 ×*g* for 5 min, then incubated for 30 min at 37 °C with 5% $CO_2$. Unbound bacteria were removed with three washes of Hanks' balanced salt solution (HBSS) washes and DMEM + 10% FBS containing 1 µg/mL Hoechst 33342 and 1:1,000 dilution of CellMask Orange. Stained cells were visualized using a Nikon Eclipse T*i*2 inverted spinning disc confocal microscope equipped with a CMOS camera (Hammamatsu ORCA-Fusion BT) and a Nikon CFI APO LWD 40X WI 1.5 NA λS objective. Live cell imaging was performed at 37°C, 5% $CO_2$ using a stage top incubator, $CO_2$ regulator, and humidity control device (OkoLabs). For each RB50 strain, 5-9 fields/well were acquired for 2 to 3 replicates. Images were processed using a combination of Fiji (https://github.com/michaelscostello/Image-analysis-scripts/tree/9936bd0744827504d1d11abd88ddb18c227789de/Fiji%20Scripts/Fiji%20pipeline%20for%20CellProfiler) and CellProfiler (https://github.com/michaelscostello/Image-analysis-scripts/tree/9936bd0744827504d1d11abd88ddb18c227789de/CellProfiler%20scripts) to segment and quantify nuclei and bacteria.

**Microtubule binding assays.** Purified α/β tubulin (450 µg) from the Microtubule Binding Protein Spin-Down Assay Biochem Kit (Cytoskeleton Inc.) was assembled into microtubules in tubulin buffer [80 mM HEPES (pH 7.0), 2 mM $MgCl_2$, 0.5 mM EGTA] supplemented with 1 mM GTP at 35 °C for 20 min. Assembled microtubules (~0.4 nM) were stabilized with 20 µM taxol. Purified FhaB-CT proteins (~5 µM) were diluted into tubulin buffer with or without stabilized microtubules and incubated at ambient temperature for 30 min. Binding reactions were layered onto 100 µL cushions of tubulin buffer containing 60% glycerol, and the samples centrifuged at 100,000 ×*g* for 40 min in a Sorvall RC M120 GX centrifuge using a fixed-angle

S100AT-3 rotor. Supernatant and pellet fractions were recovered for SDS-PAGE and immunoblot analysis.

**Mammalian cell culture and transfection.** K562 (ATCC CCL-243) cells were cultured and passaged in Roswell Park Memorial Institute 1640 medium supplemented with 10% fetal bovine serum (FBS) and 100 U/mL penicillin-streptomycin. U2OS (HTB-96) and HEK293 cells were cultured in Dulbecco's modified Eagle's media supplemented with 10% FBS and 100 U/mL penicillin-streptomycin. Vero (ATCC CCL-81) cells were a gift from Dr. Carolina Arias. Media supplemented with FBS and penicillin-streptomycin are henceforth referred to as "complete". All cell lines were maintained at 37 °C with 5% $CO_2$.

Transfection particles were prepared by mixing Mirus TransIT-LT1 reagent (7.5 µL) with OptiMEM I (250 µL) at ambient temperature for 15 min. Plasmids pCMV-*gag-pol* (ΔR8.91, 1.35 µg), pCMV-VSV-G (0.2 µg) and 1.5 µg of transfer plasmids (pINDUCER or pCW57.1) were added and allowed to form complexes for 15 min. Lipid-DNA suspensions were added to HEK293 cells (at 75% confluence) in a dropwise fashion with gentle swirling. Culture supernatants containing virus were collected after 48 h and passed through 0.45 µm cellulose acetate filters. Filtered virus was supplemented with 8 µg/mL polybrene and used to transduce K562 and U2OS cells at 60% confluence in 24-well plates. Transduced U2OS cells were enriched by selection with 1.0 µg $mL^{-1}$ puromycin. Stable integrants of transduced K562 cells were isolated by fluorescence activated cell sorting (FACS) using Sony MA900 cell sorter. Sorted cells were expanded and subjected to analytical flow cytometry to validate enrichment. Pseudoclonal populations were induced with 2 µg/mL doxycycline for 18 h, washed once with PBS, then resuspended in complete media lacking penicillin/streptomycin for infections with RB50 bacteria.

**RNA isolation and analysis.** Total RNA was extracted from K562 and K562-bacteria mixtures with 1.0 mL of 40% saturated phenol, 0.8 M guanidine isothiocyanate, 0.4 ammonium isothiocyanate, 6% glacial acetic acid, 5% glycerol (GITC-phenol). Adherent Vero cells ($1 \times 10^6$ per well) were disrupted *in situ* with 1.0 mL GITC-phenol. Total RNA was precipitated with an equal volume of 100% isopropanol, then dissolved in 10 mM Bis-tris (pH 6.5), 0.1 mM EDTA for quantification by absorbance at 260 nm. RNA samples (0.5 µg) were run on 8 M urea/1× Tris-borate-EDTA (TBE) 6% polyacrylamide gels and imaged by staining with SYBR II (Thermo Fisher Scientific). For Northern hybridization, gels were electroblotted onto Nytran$^+$ nylon membranes as described (*46*). Oligonucleotides CH6221, CH6303 and CH6305 were end-labeled with ATP-γ-[$^{32}$P] and phage T4 polynucleotide kinase and used as probes to detect 18S rRNA. Blots were visualized using a Bio-Rad Molecular Imager FX with Quantity One software.

**Ectopic expression of FhaB-CT.** U2OS cells ($5 \times 10^4$) were seeded in 24-well plates on cover glasses and grown to 50% confluence. eGFP-FhaB-CT production was induced with 1 µg/mL doxycycline for 24 h. Cells were fixed with 4% paraformaldehyde, permeabilized with PBS containing 0.5% BSA, 0.05% saponin, 50 mM $NH_4Cl$, 0.02% $NaN_3$, then incubated with a 1:250 dilution of rabbit antibodies to β-tubulin (Cell Signaling, 2146) for 1 h. Cells were washed with PBS and incubated with a 1:500 dilution of anti-rabbit secondary Star Red antibodies (Abberior, STRED-1002-500UG) for 30 min. Before visualization, cells were stained with 1 µg/mL Hoechst 33342. Coverslips were mounted onto slides using 33% w/v Mowil 4-88, 33% glycerol in PBS. Images were acquired using a Nikon Eclipse T*i*2 inverted spinning disc confocal microscope equipped with a CMOS camera (Hammamatsu ORCA-Fusion BT) and a Plan Apochromat TIRF 100×, NA 1.49 oil-immersion objective. FIJI (v2.3) (*47*) was used to process all microscopy images.

**Electron cryotomography.** *B. bronchiseptica* RB50 mini-cells were produced by *ftsZAQ* overexpression in strains CH1195 (*fhaB*$^+$) and CH1271 (Δ*fhaB*). Bacteria were grown in Steiner-Scholte medium supplemented with 5 g/L casamino acids, 1 g/L heptakis (2,6-di-*O*-dimethyl)-β-cyclodextrin and 15 µg/mL chloramphenicol at 37 °C to $OD_{600}$ of 0.2, then induced with 10 mM L-arabinose to produce mini-cells for 48 h. Bacteria were removed by centrifugation at 3,000 ×*g* for 20 min at 4 °C and the mini-cell containing supernatant harvested. Mini-cells were sedimented at 42,000 ×*g* for 1 h at 4 °C, resuspended in 10 mL of PBS, then overlaid on a discontinuous 5-20% OptiPrep gradient (Serumwerk Bernburg, Germany) for centrifugation at 4,600 ×*g* for 30 min at 4 °C. Isolated mini-cells were washed twice with PBS by repeated centrifugation steps at 42,000 ×*g* for 1 h at 4 °C, then resuspended in 100 µL of PBS for cryotomography. Aliquots (3 µL) were applied to glow-discharged Quantifoil R 2/1 Cu 200 Mesh holey carbon grids (Quantifoil, SPT Labtech, UK) and plunge-frozen into liquid ethane using a Leica EM GP2 automatic plunge freezer (Leica, Germany). Grids were imaged on a 200 kV Talos Arctica cryo-electron microscope (Thermo Fisher, USA) equipped with a Falcon 3 direct electron detector.

**X-ray crystallography.** *E. coli* CH2016 cells harboring plasmid pCH1166 were grown to $OD_{600}$ ~ 0.6 at 37 °C in LB medium supplemented with Amp. The culture was shifted to 18 °C for overnight induction with 0.5 mM IPTG. Bacteria were harvested by centrifugation at 3,000 ×*g* for 10 min, then resuspended in Buffer A [50 mM Tris-HCl (pH 7.5), 300 mM NaCl, 10% glycerol] supplemented with 0.1 mg/mL lysozyme and 1 mM phenylmethysulfonyl fluoride. After sonication, cell debris was removed by centrifugation at 11,000 ×*g* for 1 h at 4 °C. The clarified supernatant was passed through a 0.2 µm filter and loaded onto a His-trap column at 4 °C. The column was washed with Buffer A and proteins eluted with a linear gradient 0 to 500 mM imidazole in Buffer A. Fractions containing pure FhaB-CT were dialyzed against 20 mM Tris-HCl (pH 7.5), 150 mM NaCl at 4 °C and concentrated to ~75 mg/mL for crystallization trials.

Crystallization screens were performed at 4 °C, and an initial hit was found in JCSG+ condition 25 [0.2 M sodium chloride, 0.1 M phosphate-citrate (pH 4.2), 20% PEG 8000]. To obtain phase information, selenomethionine (SeMet)-labelled FhaB-CT was produced and purified as described above (*48*). After three months, crystals were harvested from 0.2 M NaCl, 0.1 M phosphate-citrate (pH 5.0), 26.5% PEG 8000. Crystals were cryo-cooled in mother liquor supplemented by 15% glycerol and were used in X-ray diffraction experiments at ALS 5.0.2. A multiwavelength anomalous dispersion (MAD) dataset was collected at two wavelengths, 0.9794 Å and 1.0 Å. The data were indexed and integrated in xds version Jan 10, 2022 (*49*), and scaled in aimless (version 0.7.4) to 1.65 Å resolution (*50*). The structure was solved and an initial model was built in AutoSol in Phenix 1.201-4487 (*51*). Rounds of refinement were carried out in phenix.refine and Coot (versions 0.9.8.1 - 0.9.8.8) to a final $R_{work}/R_{free}$ of 17.9%/20.4% (*52, 53*). All data collection and refinement statistics are presented in **Table S1**.

**Negative-stain electron microscopy.** All samples were negatively stained as described (*54*). Samples (3 µL) at various concentrations were pipetted onto positively glow-discharged carbon-coated 200-mesh Gilder Copper grids (Ted Pella). Excess sample was removed with filter paper, washed 3 times with 50 µL Milli-Q water drops, and stained with two 50 µL drops of freshly prepared 0.75% uranyl formate (Electron Microscopy Sciences). Excess stain was vacuum aspirated. Grids were carbon-coated using a Leica ACE200 and positively glow-discharged using a PELCO easiGlow (Ted Pella). All grids were imaged using a JEOL JEM-2100F TEM equipped with a Gatan OneView 4k x 4k camera. Micrographs in **Fig. S7** were contrast enhanced using Fiji (*47*).

**Cryogenic electron microscopy.** $His_6$-FhaB-CT was purified and bound to *in vitro* assembled microtubules as described above. FhaB-CT•microtubule complexes were isolated by centrifugation at 109,000 x *g* for 1.5 h at 20 °C using a Beckman Coulter Optima L-90K Ultracentrifuge with a SW55Ti rotor. Complexes were analyzed by SDS-PAGE to assess sample quality and applied to cryo-EM grids immediately. Complexes were deposited onto Quantifoil R2/1 300 mesh Cu grids which were positively glow discharged using a PELCO easiGlow (Ted Pella). A Leica EM GP2 set to 4 °C and 95% humidity was used to freeze the grids. Sample (3 µL at 0.83 mg/mL) was applied to the front of the grid prior to front blotting for 4 s. Grids were plunge-frozen in liquid ethane and stored in liquid nitrogen.

Grids were screened using a 200 keV Talos Glacios (Thermo Fisher Scientific) to assess ice thickness, filament quality and distribution. All movies were collected with a 300 keV FEI Titan Krios microscope equipped with a Thermo Fisher Scientific Selectris X energy filter and a Falcon 4i direct electron detector. Movies were collected using SerialEM at a pixel size of 1.196 Å/pixel with a total dose of 50 $e^-/Å^2$ spread over 2673 total frames, a defocus range of -0.8 to -2 µm and a 100 µm objective aperture. The energy filter slit width was set to 6 eV. Full data collection parameters are provided in **Table S2**.

The processing pipelines are illustrated in **Figs. S8** & **S10**. CryoSPARC 4.3.1 (*55*) was used to process 20,436 movies with patch motion correction (*56*) and patch contrast transfer function (CTF) estimation. An initial set of 889 images were used to blob pick 116,007 particles to create initial 2D templates for filament tracer. After multiple rounds of 2D classification

48,633 particles were used to create an *ab initio* helical refined 3.97 Å map with an optimized helical twist of -0.609°, a helical rise of 81.984 Å and a helical order of 2. This map was used to create templates for the filament tracing of 19,222 selected images. A filament diameter of 280 Å and a separation distance between picks of 0.3 diameters or 84 Å, the distance associated to the α- and β-tubulin heterodimer, resulted in 2,197,679 extracted particles. After multiple rounds of 2D classification, 940,421 particles were selected for *ab initio* non-uniform helical refinement resulting in a 3.22 Å map (deposited under EMD-49766) with an optimized helical twist of -0.562°, a helical rise of 81.992 Å and a helical order of 1.

Individual protofilaments unbound and bound to FhaB-CT were isolated from the 3.22 Å map using atomic models of α-tubulin, β-tubulin (from PDB: 7TQY) and FhaB-CT (created using AlphaFold2 (*57*)) using the color zone feature in UCSF ChimeraX 1.7 (*58*). Reference maps of various microtubule architectures were created using the isolated protofilaments and the following maps as templates: EMD-5192 for 12-3, EMD-8997 for 13-3, EMD-8998 for 14-3 and EMD-5195 for 15-4 (*59, 60*). Reference maps were subjected to heterogeneous refinement which resulted in 11.9% for 12-3, 7.6% for 13-3, 74.5% for 14-3, and 6.0% for 15-4 of the total 1,966,917 particles. 14-3 microtubules were used for subsequent data processing.

Non-uniform helical refinement using the 14-3 reference map lowpass filtered to 35 Å resulted in a 3.10 Å map with a helical twist of -0.531°, a helical rise of 82 Å and a helical order of 1. A mask around the highest resolved 4 protofilaments by 4 monomers (2 dimers) section of the map, using the color zone feature in UCSF ChimeraX, was created for particle subtraction. The subtracted particles were subjected to 3D classification into 4 classes and 3 classes, composed of 1,193,804 particles, were selected for the final maps. Subsequent local CTF refinement, global CTF refinement and local refinement resulted in a final 2.89 Å 14-3 microtubule reconstruction (deposited under EMD-49767). Another round of masking and particle subtraction using the 2.89 Å map resulted in a 2.52 Å focused FhaB-CT bound microtubule map. Subsequent local CTF refinement, global CTF refinement and local refinement resulted in a final 2.42 Å focused FhaB-CT bound microtubule reconstruction (deposited under EMD-49577 and PDB:9NNL). A similar process was employed to mask the region around the seam for particle subtraction. The subtracted particles were subjected to 3D classification into 4 classes and a single class, composed of 169,390 particles, was selected for the final maps. Subsequent processing resulted in a final 3.16 Å aligned 14-3 microtubule reconstruction (deposited under EMD-49768) and a final 2.93 Å focused microtubule seam reconstruction (deposited under EMD-49769).

Atomic models for α- and β-tubulin were adapted from PDB:7TQY, and the FhaB-CT model was generated with AlphaFold2 (*57*). An atomic model for a 2 protofilaments by 3 monomers patch was rigid body fit with UCSF ChimeraX and positioned to best fit the cryo-EM density map using *Coot* 0.9.8.95 (*61*). The model was refined into the final map using *Phenix* real-space refinement v1.21.2-5419 (*62, 63*). Model validation was performed using *Phenix* and MolProbity (*64*). Statistics are available in **Table S2**.

**Trachea explant infections and microscopy.** Sprague-Dawley rats were housed in the animal resource center at the University of California Santa Barbara using a 12 h light/dark cycle with *ad libitum* access to food, water and enrichment. All animal studies were performed with the approval of the University of California Santa Barbara Institutional Animal Care and Use Committee. Animals were weaned at day 21, separated by sex, group-housed and randomly assorted. Experimental replicates from different litters were used for analyses. Male rats (8 - 12 w) were anesthetized with ketamine (200 mg/kg) and xylazine (20 mg/kg) in saline then sacrificed by cervical dislocation. Tracheas were exposed with a vertical incision from the base of the sternum to the mandible and excised after separation of the esophagus and surrounding vasculature. Tracheas were flushed with complete DMEM using a 20-gauge needle and syringe to remove air bubbles and blood before being placed into ice-cold complete DMEM. Connective tissue and vasculature were removed from tracheas using a #22 scalpel on a 3D printed cutting board (https://github.com/michaelscostello/TracheaTools/blob/0ff65f9dc67bcc3fe7389e3a94163de2a0ff56f4/Trachea_CB_V3.STL), and transverse incisions were made between cartilage rings to produce tracheal rings. Rings from individual animals were washed once in complete DMEM, then placed together in a 6-well plate containing 4 mL complete DMEM for incubation at 37 °C with 5% $CO_2$. For live imaging experiments, tracheal rings were placed in glass bottom 96-well plates and infected with 3 x $10^7$ cfu of RB50 in antibiotic-free DMEM for 30 min. Tracheal rings were washed three times with 200 µL Hank's balanced salt solution (HBSS) and placed into 3D-printed scaffolds (https://github.com/michaelscostello/TracheaTools/blob/0ff65f9dc67bcc3fe7389e3a94163de2a0ff56f4/Trachea_SC_V14.STL) to immobilize the tissue samples. Explants were incubated with Tubulin Tracker™ Deep Red (1 µM) and Hoechst 33342 (1 µg/mL) and bacterial-epithelial interactions visualized using a spinning disc microscope (Nikon Eclipse T*i*2 equipped with a Yokogawa spinning disc unit an Orca Fusion BT ssMOS camera) using a CFI APO LWD 40X WI 1.5 NA λS objective. All imaging was performed at 37°C, 5% $CO_2$ using a stage top incubator, $CO_2$ regulator, and humidity control device (OkoLabs). Image series were acquired with NIS Elements (Nikon), and included a combination of z-stacks and time-lapse series.

Z-stack series were cropped to produce unique fields containing patches of confluent ciliated epithelia in Fiji v2.3. For each cilia patch, a line was drawn at the base of cilia in the 647 nm channel. Line coordinates were determined using the FIJI script “Get_line_coordinates.ijm” (https://github.com/michaelscostello/TracheaTools/blob/0ff65f9dc67bcc3fe7389e3a94163de2a0ff56f4/Get_line_coordinates.ijm) and recorded in a spreadsheet. Bacterial centroids were determined the 568 nm channel using the FIJI script "Centroid.ijm” (https://github.com/michaelscostello/TracheaTools/blob/0ff65f9dc67bcc3fe7389e3a94163de2a0ff56f4/Centroid.ijm) and recorded in a spreadsheet. The Euclidian distance between the line at the cilia base and bacterial centroid was determined using the excel macro “Closesest_Distance()" (https://github.com/michaelscostello/TracheaTools/blob/0ff65f9dc67bcc3fe7389e3a94163de2a0ff56f4/Closest_Disance().rtf). Individual distance values for each bacterium were plotted as dots, with each color representing a biological replicate and colored diamonds representing the mean distance from the cilia base of each biological replicate.

**Mouse nasal colonization.** Mouse respiratory colonization was performed with *B. pertussis* Tohama I (CIP 81.32 as described previously (*9, 65*). Bacteria for intranasal infections were grown for 48 h on nutritionally enriched BG plates containing modified Stainer–Scholte medium supplemented with 3 g/L casamino acids. Bacteria were harvested from the agar and suspended in prewarmed PBS for intranasal inoculation. In-house bred C57BL/6J MyD88$^{-/-}$ female and male mice (MyD88 null, B6.129P2(SJL)-Myd88tm1.1Defr/J, Jackson Laboratory, USA) were anesthetized by intraperitoneal injection of ketamine (80 mg/kg) and xylazine (8 mg/kg) in 0.9 % saline and challenged intranasally with 5 µL of $1\times10^7$ cfu of Tohama I strains. At 7 d, mice were sacrificed and the dissected nasal cavities with turbinates were homogenized in PBS using an IKA Ultra Turrax T25 tissue homogenizer (Sigma-Aldrich, USA). Suspensions were cleared of bone debris by centrifugation at 217 ×*g* for 10 min, and serial dilutions of the supernatant were plated onto BG agar supplemented with 15% defibrinated sheep blood and 100 µg/mL streptomycin. Bacterial cfu were enumerated after 4 days at 37 °C.

All animal experiments were approved by the Animal Welfare Committee of the Institute of Molecular Genetics of the Czech Academy of Sciences, Prague, Czech Republic. Handling of animals was performed according to the Guidelines for the Care and Use of Laboratory Animals, the Act of the Czech National Assembly, Collection of Laws no. 246/1992. Permissions no. 1989/2023 was issued by the Animal Welfare Committee of the Institute of Molecular Genetics, Czech Academy of Sciences, Prague, Czech Republic.

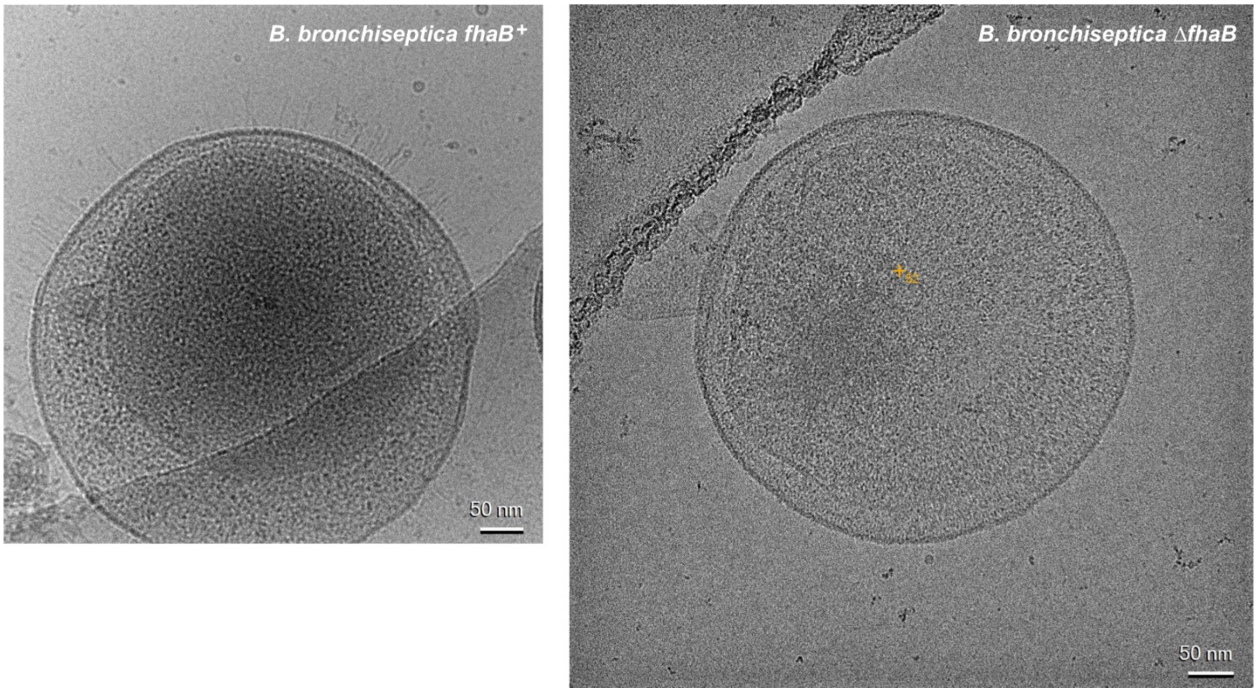

**Fig. S1. Electron cryotomography of *B. bronchiseptica* RB50 mini-cells.** Mini-cells derived from RB50 strains CH1195 (Δ*fim fhaB*$^{+}$) and CH1271 (Δ*fim* Δ*fhaB*) were image by electron cryotomography as described in Materials and Methods.

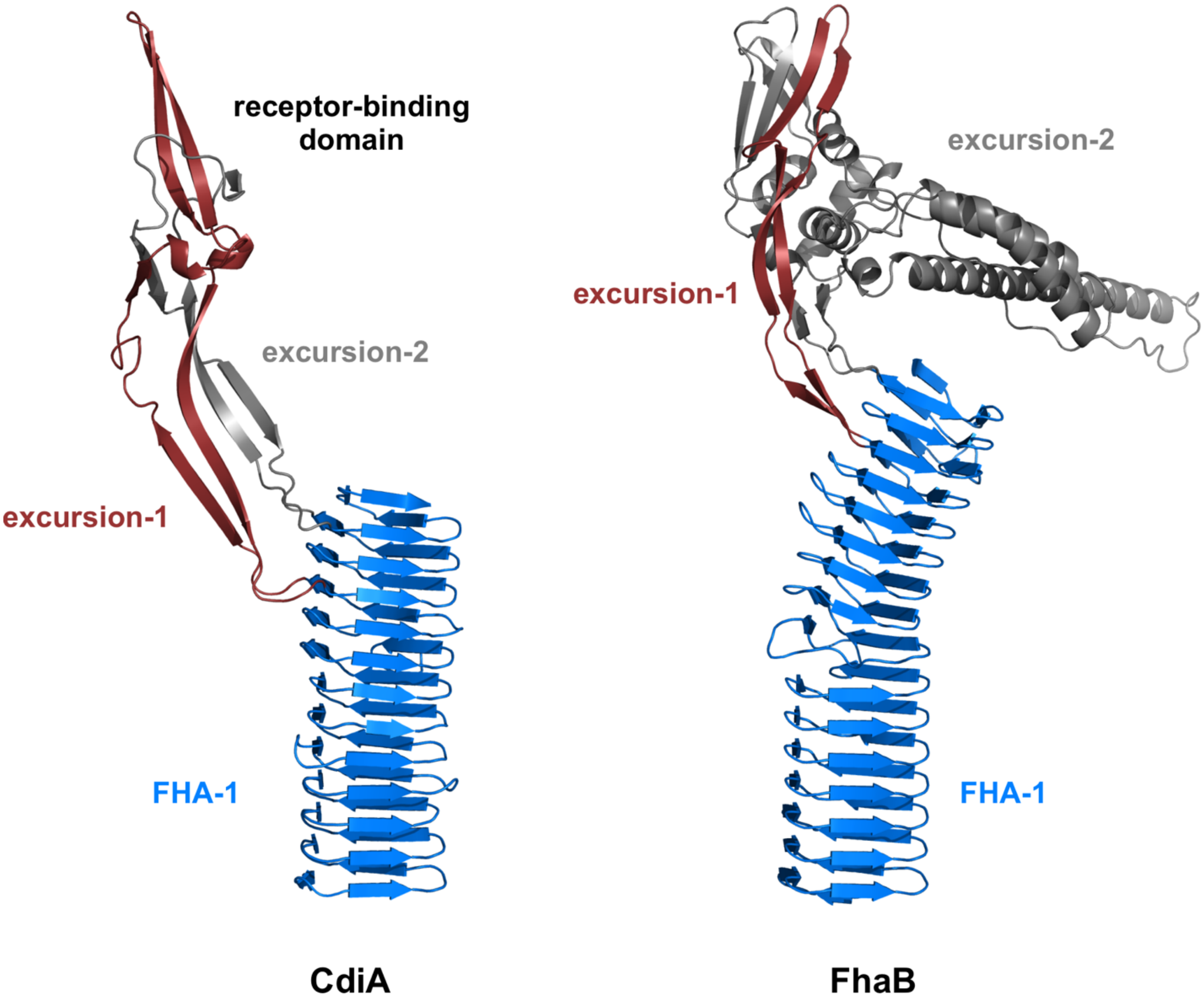

**Fig. S2. AlphaFold3 models of receptor-binding domains.** The receptor-binding regions of CdiA (GenBank: EJK94116.1) and FhaB (NCBI: WP_041936401.1) were modeled using AlphaFold3. Predicted excursion-1 and excursion-2 subdomains are colored in brick red and gray, respectively.

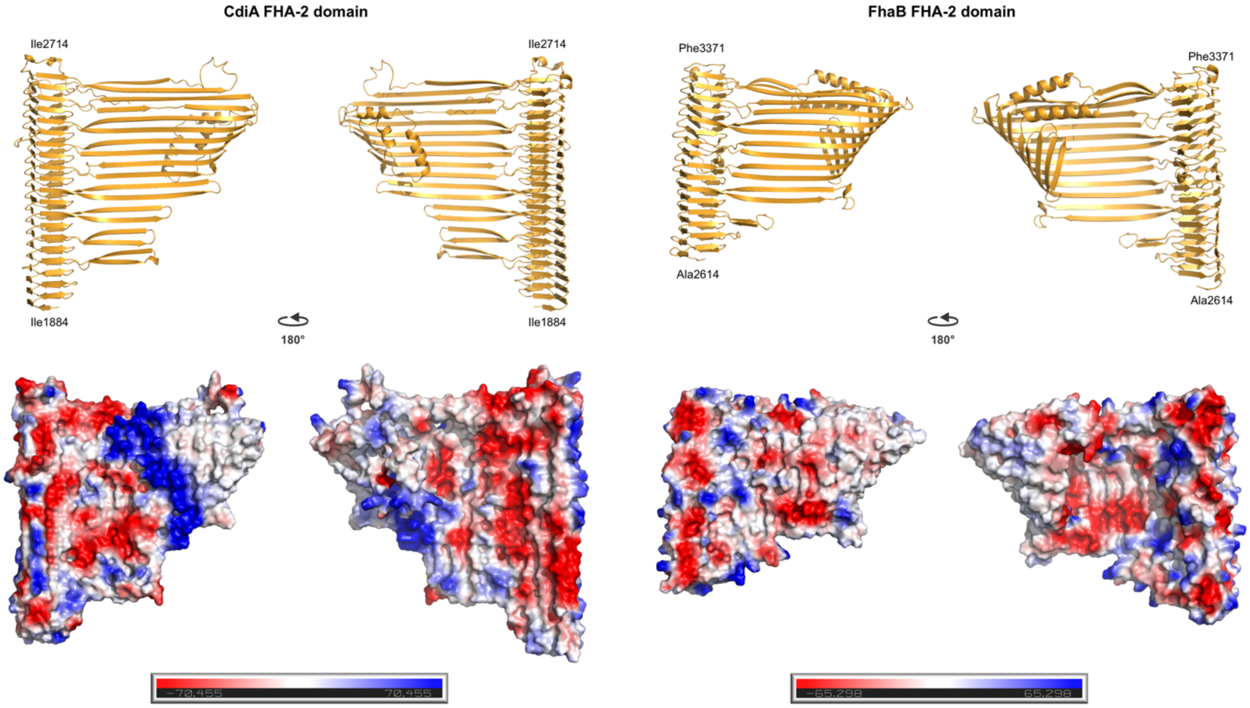

**Fig. S3. AlphaFold3 models of FHA-2 domains.** The FHA-2 domains of CdiA from *E. coli* STEC_O31 (GenBank: EJK94116.1) and FhaB from *B. bronchiseptica* RB50 (NCBI Ref Seq: WP_041936401.1) were modeled using AlphaFold3. Predicted structures are rendered as cartoons and as electrostatic surface potential maps.

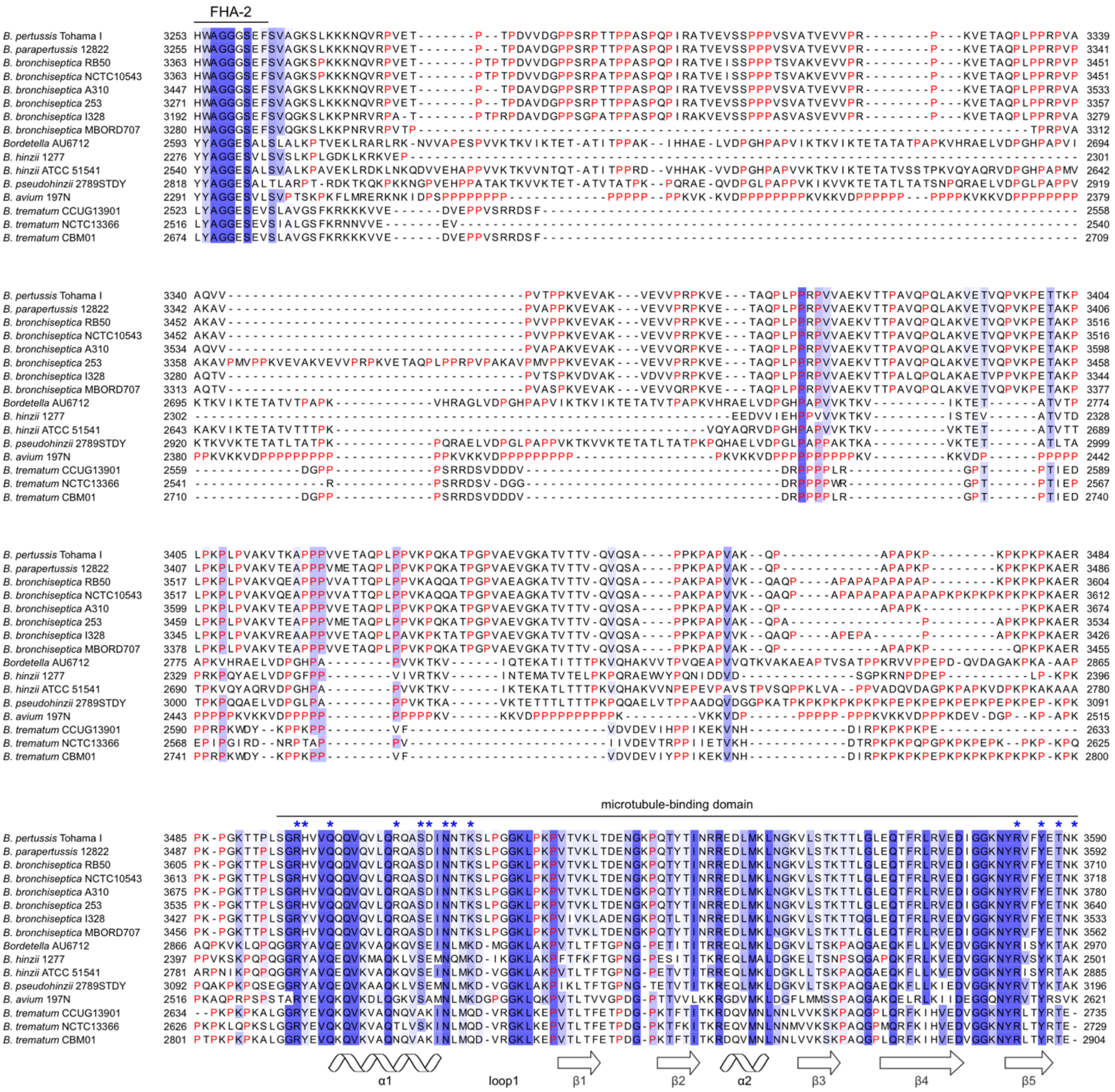

**Fig. S4. Alignment of proline-rich and CT domains of *Bordetella* FhaB proteins.** FhaB proteins from *B. pertussis* Tohama I (NCBI: WP_010930610.1), *B. parapertussis* 12822 (WP_010928856.1), *B. bronchiseptica* RB50 (WP_041936401.1), *B. bronchiseptica* NCTC 10543 (WP_126621964.1), *B. bronchiseptica* A310 (WP_101556242.1), *B. bronchiseptica* 253 (CCJ52713.1), *B. bronchiseptica* I328 (WP_068925961.1), *B. bronchiseptica* MBORD707 (WP_050483240.1), *Bordetella* AU6712 (WP_244193841.1), *B. hinzii* 1277 (WP_155271618.1), *B. hinzii* ATCC 51541 (AHV94525.1), *B. pseudohinzii* 2789STDY (WP_109842424.1), *B. avium* 197N (WP_012417632.1), *B. trematum* CCUG13901 (WP_025516388.1), *B. trematum* NCTC 13366 (WP_125117792.1) and *B. trematum* CBM01 (QIM72507.1) were aligned with Clustal

Omega. The alignment was rendered with Jalview at 30% sequence identity. Proline residues are depicted in red font. RB50 FhaB-CT residues that make contact with microtubules are indicated by blue asterisks (*) above the alignment, and secondary structure elements are shown below the alignment.

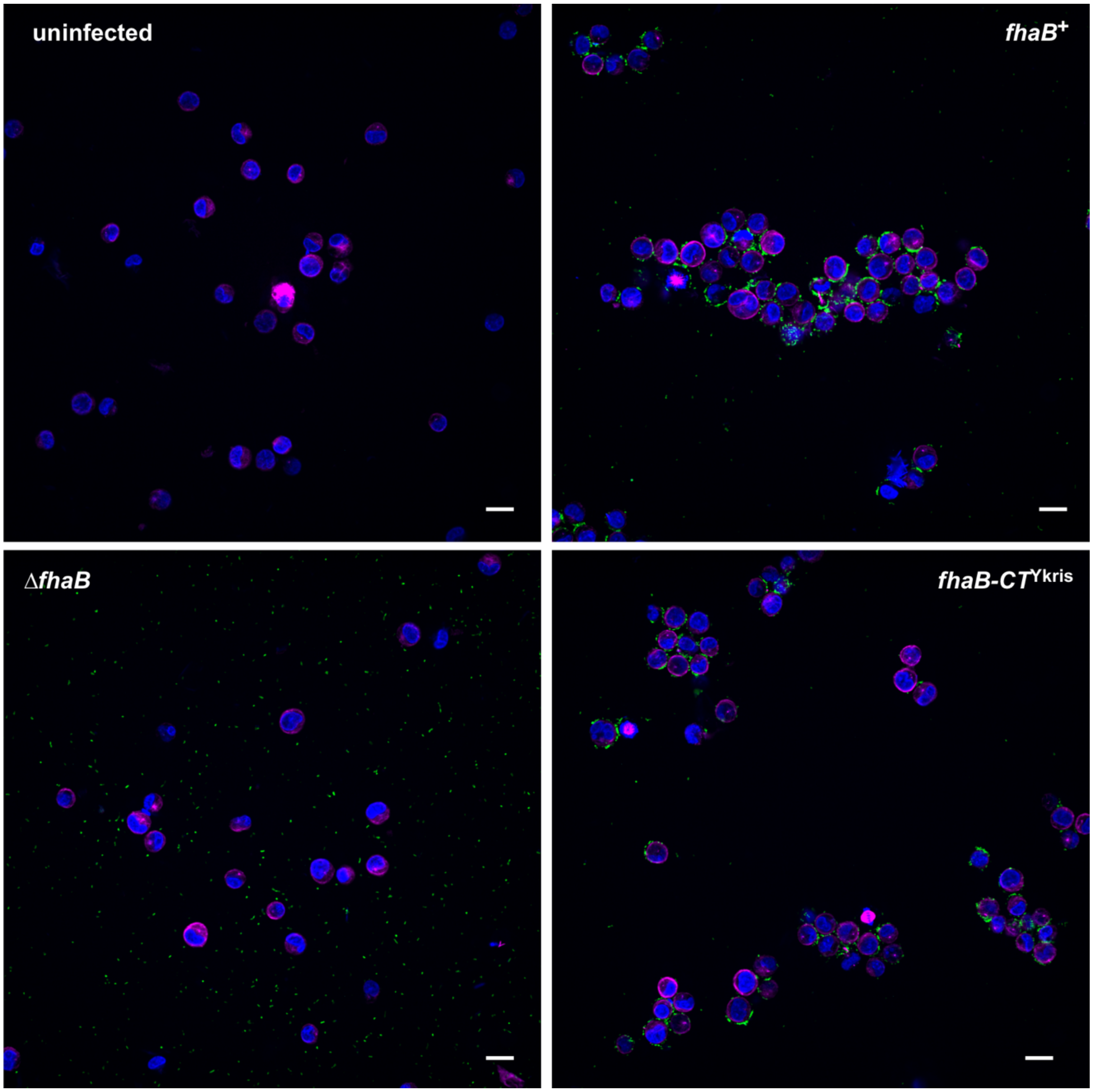

**Fig. S5. FhaB and FhaB-CT$^{Ykris}$ agglutinate K562 cells.** K562 cells were centrifuged with the indicated mNeon labeled *B. bronchiseptica* RB50 strains. After incubation for 10 min, the cell suspensions were labeled with tubulin-tracker and Hoeschst dyes to visualize microtubules and cell nuclei, respectively. Scale bars = 10 µm.

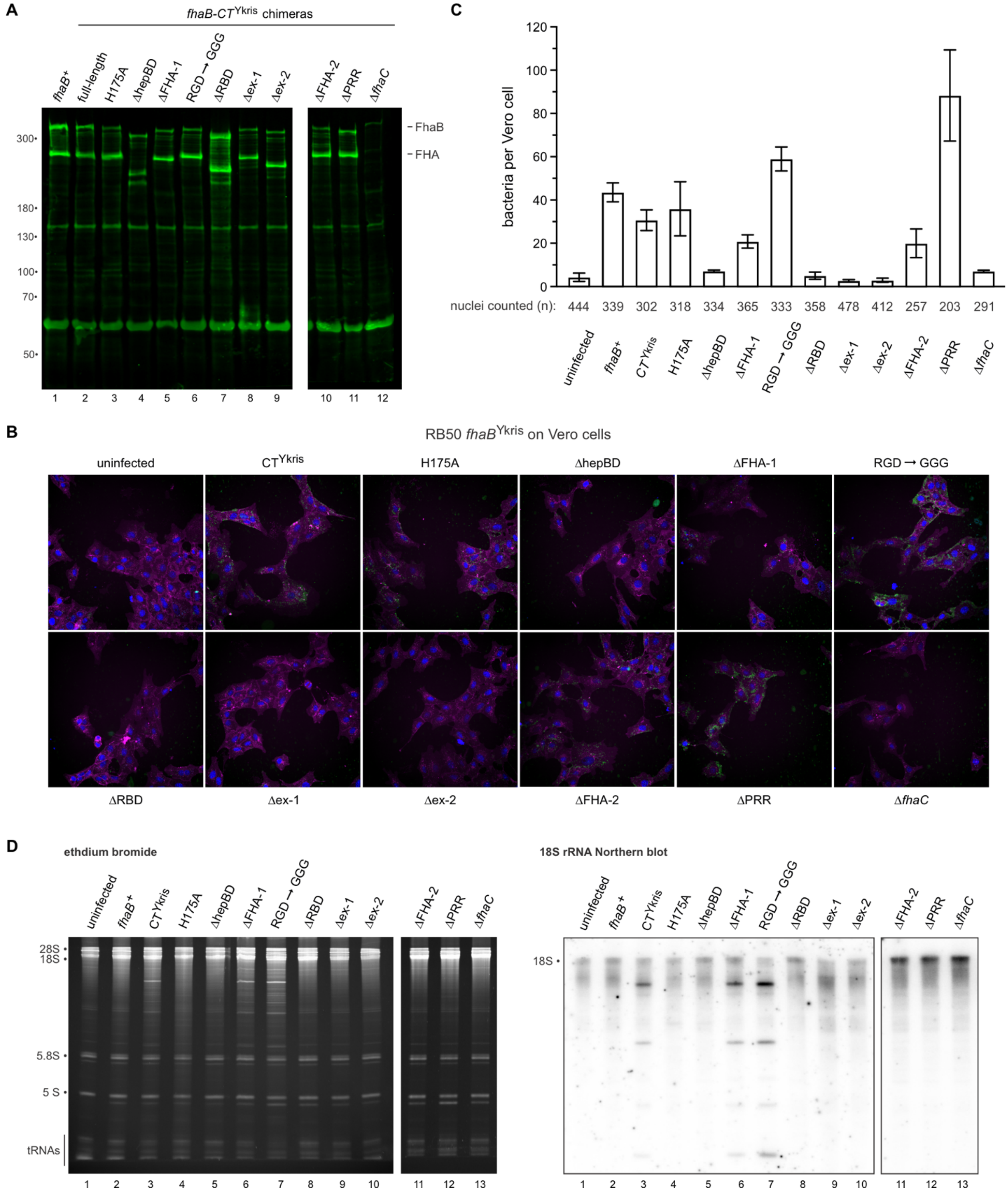

**Fig. S6. FhaB-CT$^{Ykris}$ mediated adhesion and RNase delivery to Vero cells.** (**A**) Immunoblot of FhaB-CT$^{Ykris}$ chimeras using polyclonal antibodies the N-terminal TPS transport domain of FhaB. (**B**) Representative micrographs of mNeon-labeled *B. bronchiseptica* bound to Vero cells. (**C**) Quantification of bacterial adhesion to Vero cells. Fiji and CellProfiler were used to segment and quantify cell nuclei and bacteria as described in Methods. Data are presented as averages ± SEM for the indicated number (*n*) of enumerated nuclei. (**D**) Northern blot analysis of 18S rRNA cleavage mediated by FhaB-CT$^{Ykris}$ chimeras.

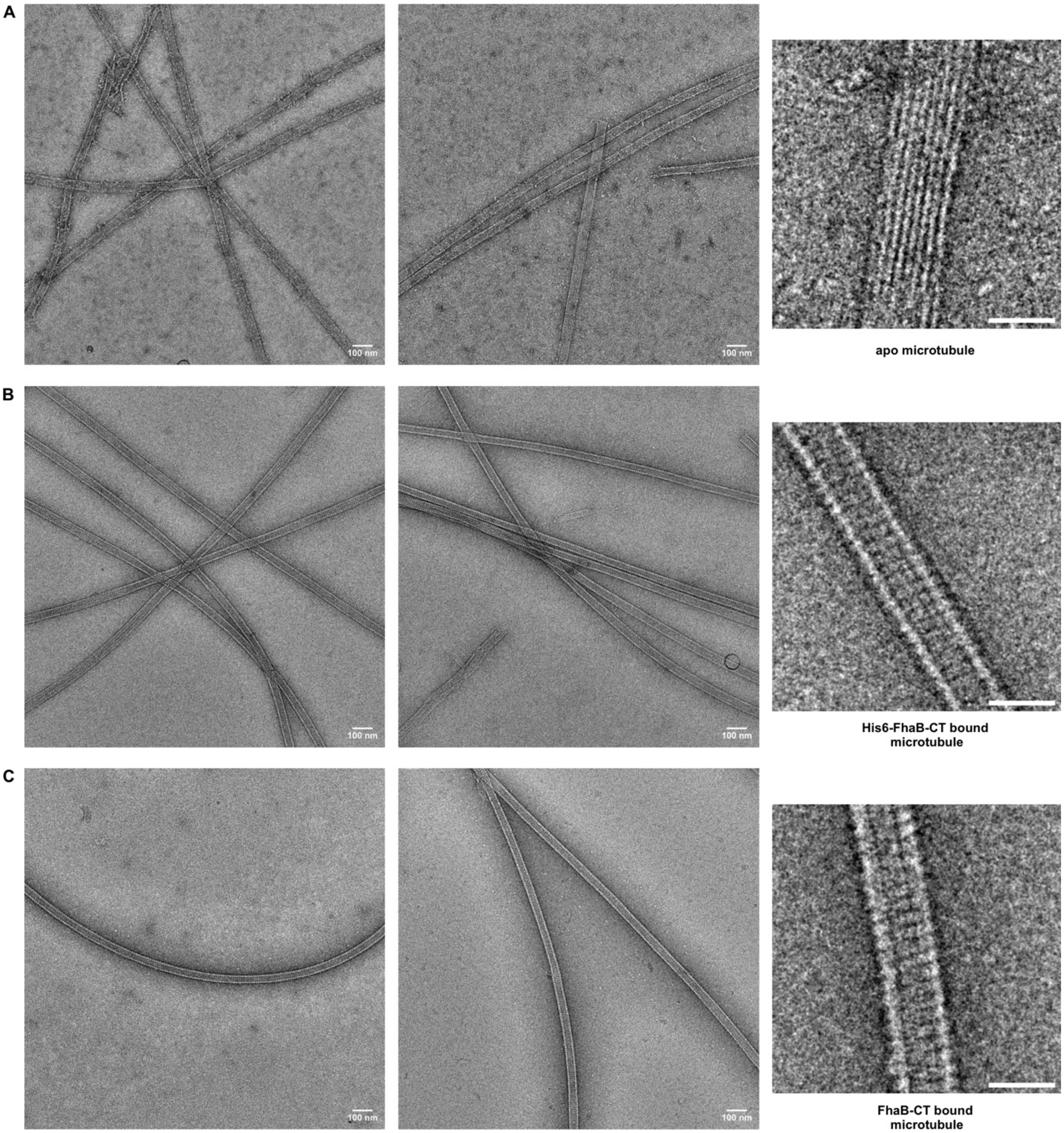

**Fig. S7. Negative-stained electron microscopy images of microtubules.** (**A**) Assembled microtubules without FhaB-CT (**B**) $His_6$-FhaB-CT bound to *in vitro* assembled microtubules. (**C**) Untagged FhaB-CT bound to *in vitro* assembled microtubules. Zoomed images, scale bar = 50 nm.

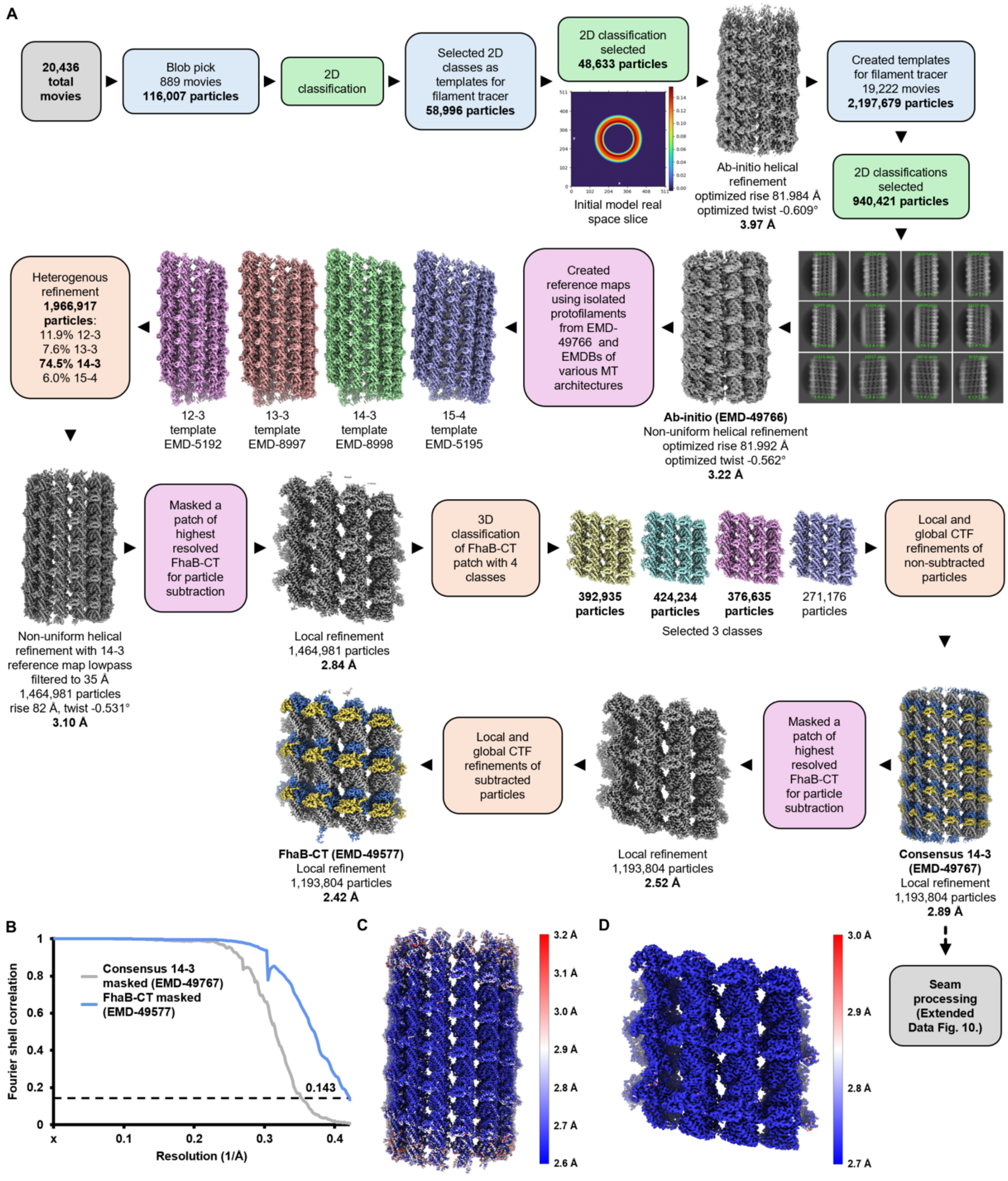

**Fig. S8. Processing flowchart for FhaB-CT using CryoSPARC.** (**A**) Processing workflow resulting in the consensus 14-3 and FhaB-CT reconstructions. (**B**) Fourier shell correlation (FSC) plot of the two final reconstructions. FSC = 0.143 is indicated by a dashed black line. Local resolution of (**C**) consensus 14-3 and (**D**) FhaB-CT calculated within CryoSPARC.

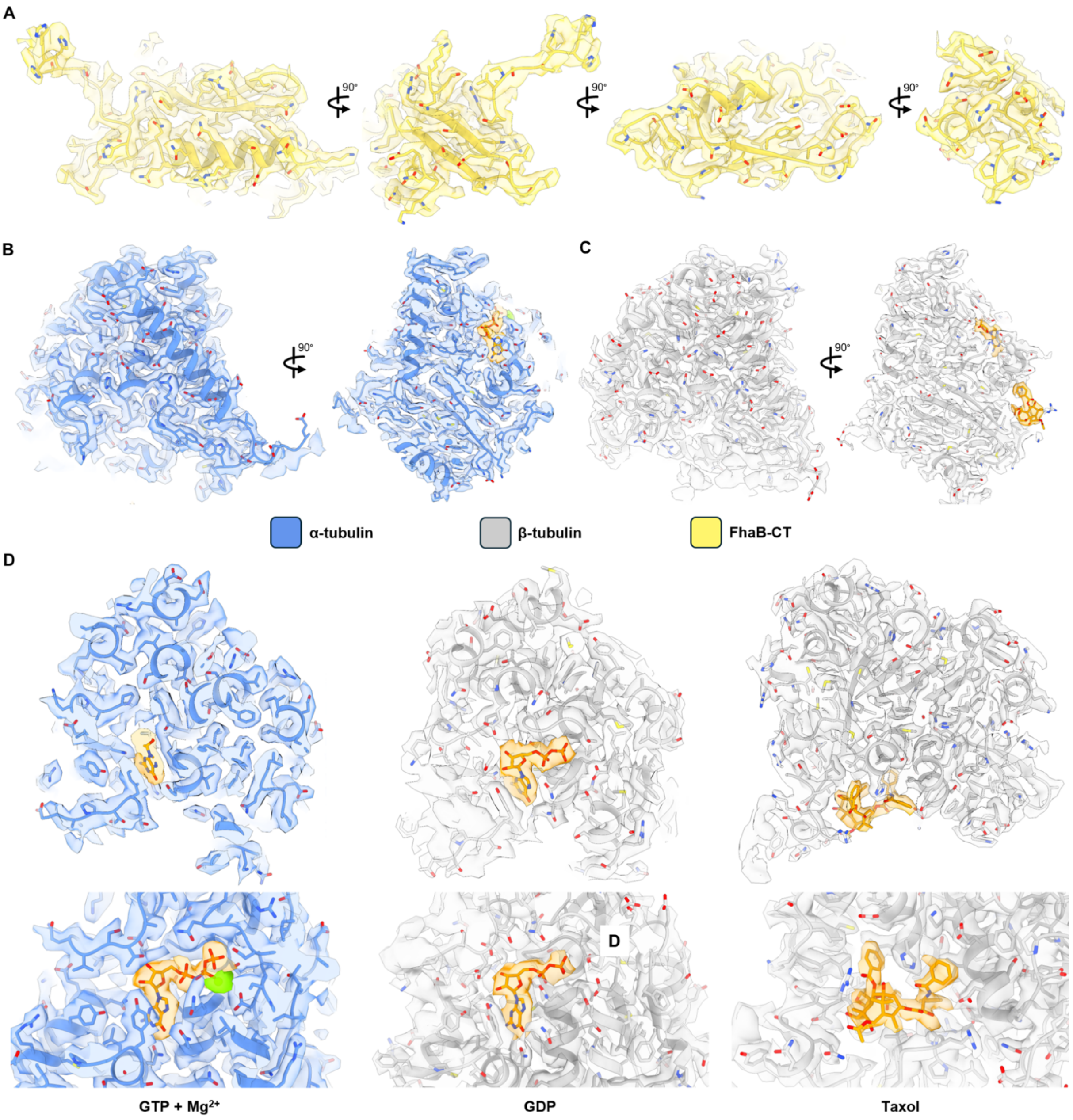

**Fig. S9. Model-to-map fits of structural features.** (**A**) FhaB-CT, (**B**) α-tubulin, (**C**) β-tubulin, and (**D**) indicate ligands fitted into the FhaB-CT map (EMD-49577).

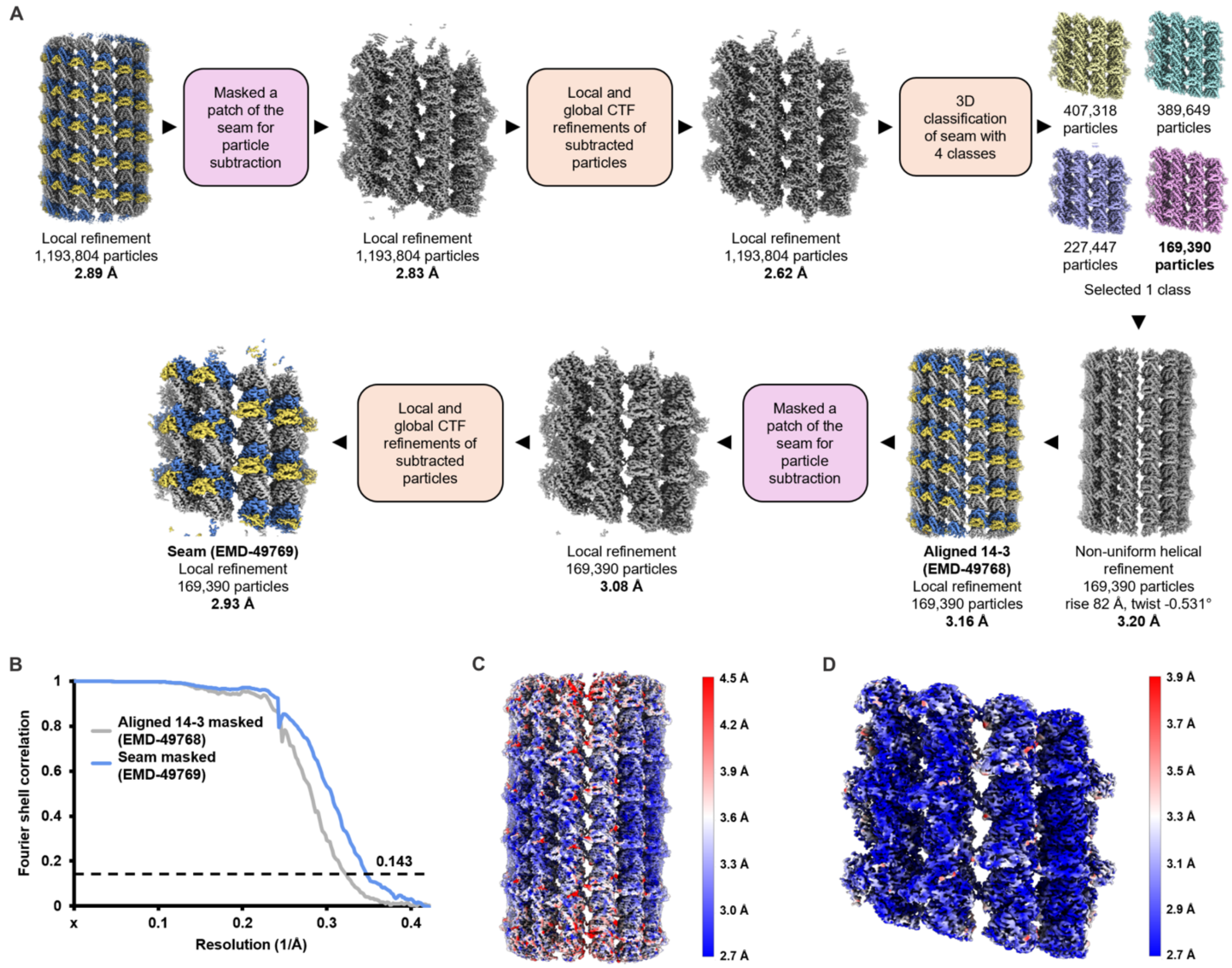

**Fig. S10. Processing flowchart for the seam using CryoSPARC.** (**A**) Processing workflow resulting in the aligned 14-3 and the seam reconstructions. (**B**) Fourier shell correlation (FSC) plot of the two final reconstructions. The dashed black line indicates FSC = 0.143. Local resolution of (**C**) the aligned 14-3 and (**D**) the seam calculated within CryoSPARC.

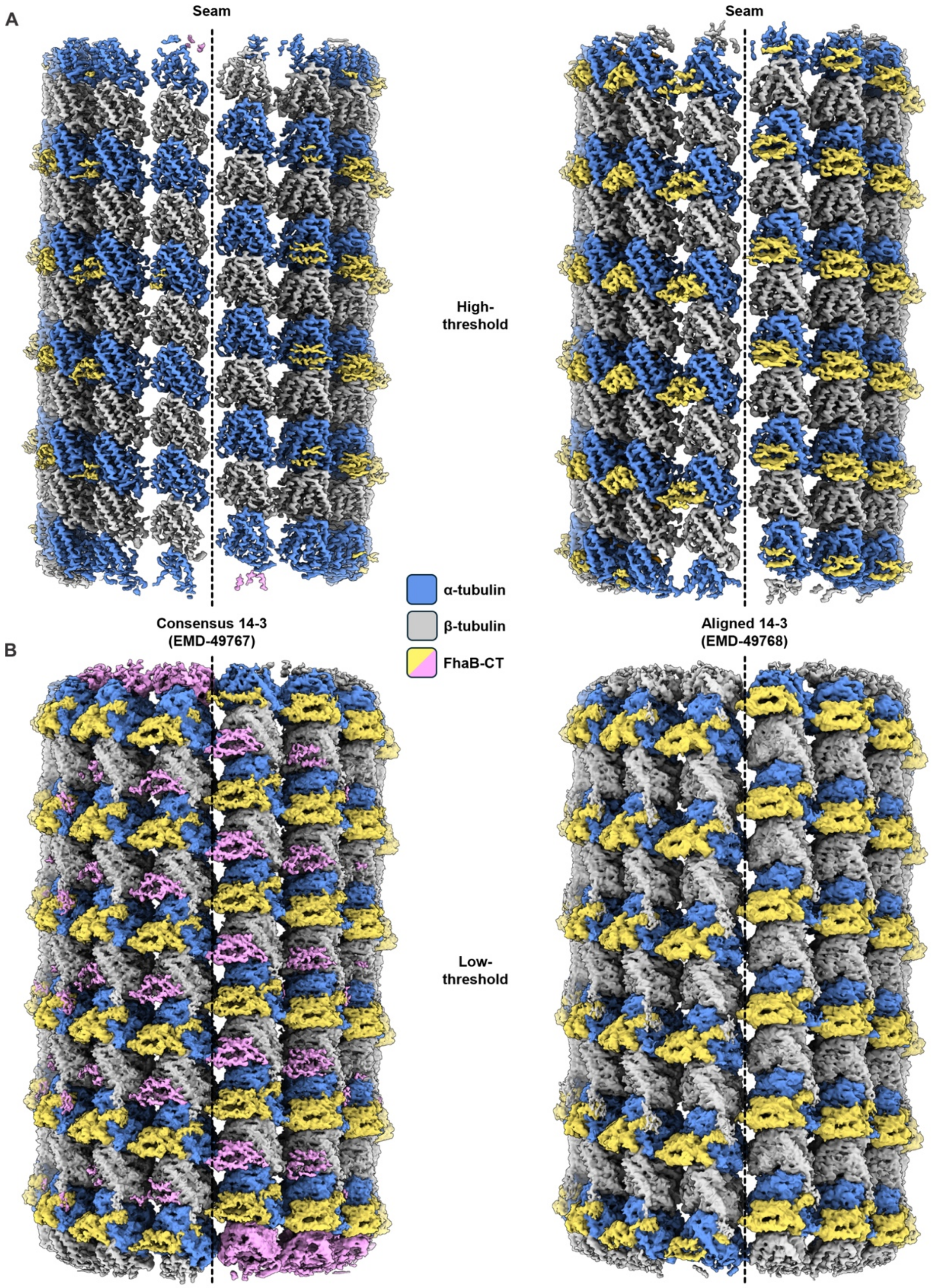

**Fig. S11. FhaB-CT binds to the microtubule seam.** (**A**) High-threshold and (**B**) low-threshold of the consensus 14-3 (left) or aligned 14-3 (right) highlighting FhaB-CT at the microtubule seam.

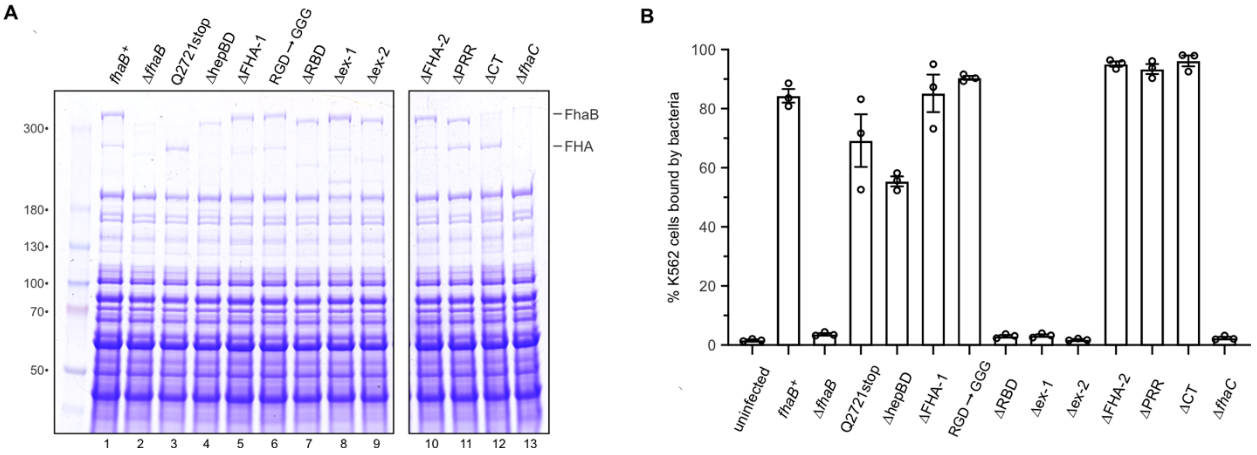

**Fig. S12. FhaB variant expression and adhesion to K562 cells.** (**A**) SDS-PAGE analysis of FhaB variants. (**B**) Bacterial adhesion to K562 cells. The percentage of K562 bound to fluorescent bacteria was quantified by flow cytometric counting of 10,000 events. Data are presented as averages ± SEM for three independent experiments.

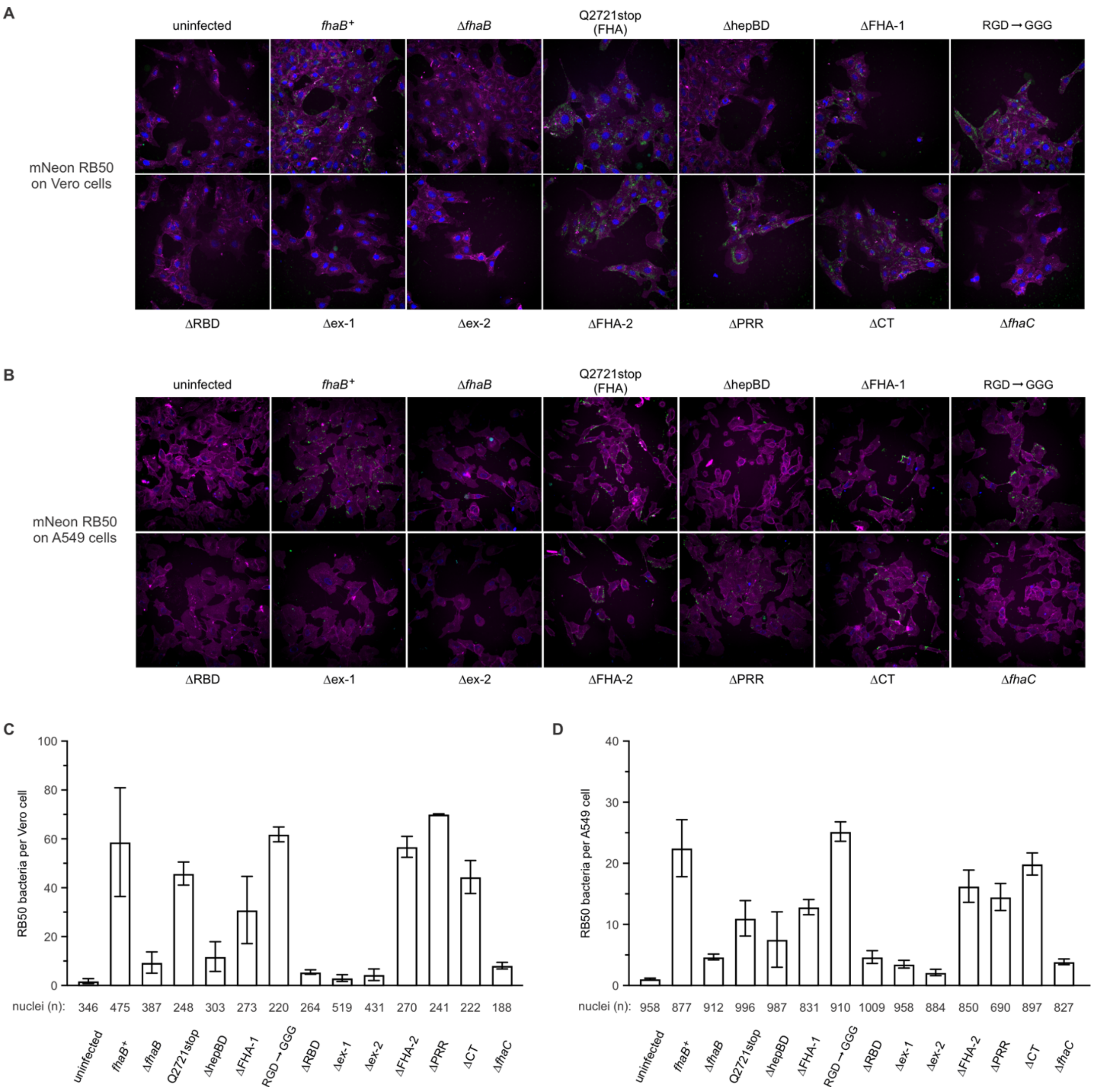

**Fig. S13. FhaB mediated adhesion Vero and A549 cells.** (**A**) Representative micrographs of mNeon-labeled *B. bronchiseptica* bacteria bound to Vero cells. (**B**) Representative micrographs of mNeon-labeled *B. bronchiseptica* bacteria bound to A549 cells. Quantification of bacterial adhesion to Vero (**C**) and A549 (**D**) cells. Fiji and CellProfiler were used to segment and quantify cell nuclei and bacteria as described in Methods. Data are presented as averages ± SEM for the indicated number (*n*) of enumerated nuclei.

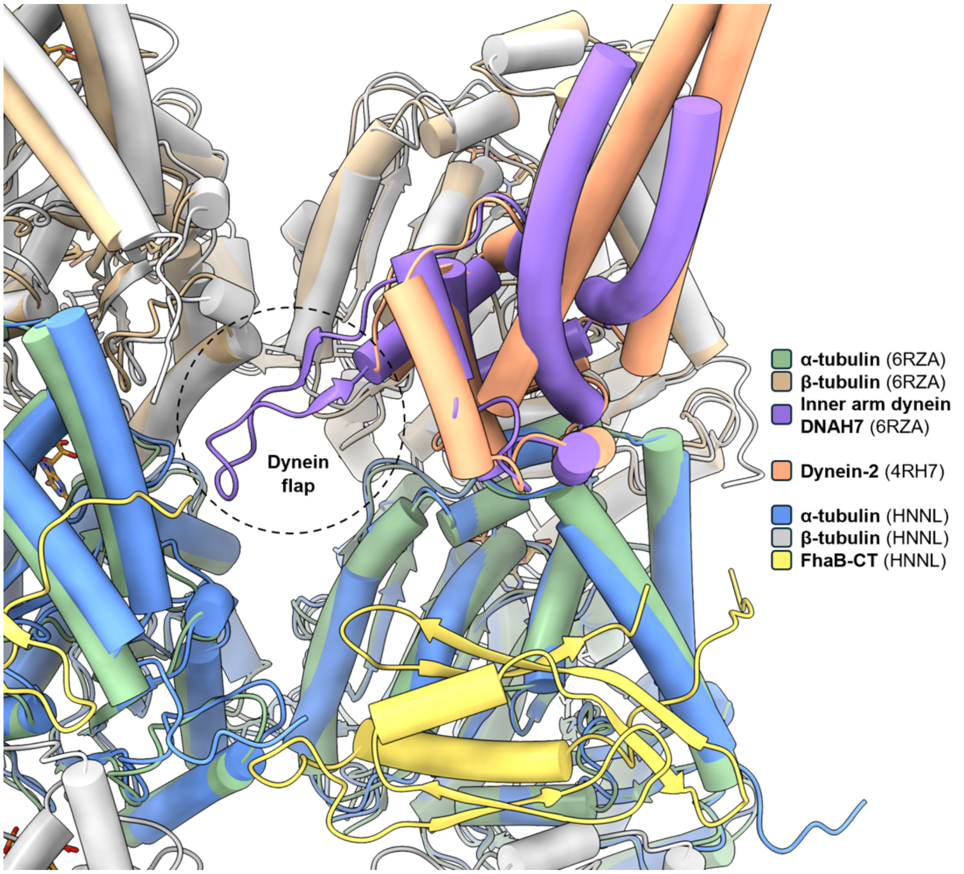

**Fig. S14. Binding site of FhaB-CT compared to dynein microtubule-binding sites.** Binding of the inner arm dynein DNAH7 (PDB: 6RZA) to the βV binding site. Both DNAH7 and FhaB-CT contact the adjacent protofilament to possibly stabilize and induce flattening of the microtubule, whereas the retrograde transporter dynein-2 (PDB: 4RH7) does not possess a dynein flap.

**Table S1. Crystallography data collection and refinement statistics.**

| | SeMet-FhaB-CT | |
|---|---|---|
| **Data collection** | | |
| Space group | $P\ 2_1$ | |
| Cell dimensions | | |
| *a*, *b*, *c* (Å) | 23.1, 46.6, 76.0 | |
| α, β, γ (°) | 90, 91.7, 90 | |
| | *peak* | *native* |
| Wavelength (Å) | 0.9794 | 1.0 |
| Resolution (Å) | 46.58-1.65 (1.68-1.65) [a] | 46.5-1.58 (1.61-1.58) |
| $R_{merge}$ [b] | 0.060 (0.805) | 0.049 (0.604) |
| $CC_{1/2}$ | 0.999 (0.704) | 0.999 (0.675) |
| $I / \sigma I$ | 15.0 (2.2) | 14.6 (2.2) |
| Completeness (%) | 98.8 (99.6) | 98.2 (97.7) |
| Redundancy | 6.7 (6.9) | 4.7 (4.4) |
| **Refinement** | | |
| Resolution (Å) | 39.71-1.65 (1.71-1.65) | |
| No. reflections | 19249 (1891) | |
| $R_{work}$ / $R_{free}$ [c] | 17.9/20.4 | |
| Ramachandran favored (%) | 98.9 | |
| Ramachandran outliers (%) | 0 | |
| No. atoms | | |
| Protein | 1587 | |
| Ligand/ion | 31 | |
| Water | 125 | |
| *B*-factors | | |
| Protein | 28.7 | |
| Ligand/ion | 45.7 | |
| Water | 36.5 | |
| R.m.s deviations | | |
| Bond lengths (Å) | 0.009 | |
| Bond angles (°) | 1.080 | |
| PDB identifier | 8SXO | |

[a] Values within parentheses refer to the highest resolution shell.

[b] $R_{merge} = \sum\sum|I_{hkl} — I_{hkl}(j)|/\sum I_{hkl}$, where $I_{hkl}(j)$ is observed intensity and $I_{hkl}$ is the final average value of intensity.

[c] $R_{work} = \sum||F_{obs}| - |F_{calc}||/\sum|F_{obs}|$ and $R_{free} = \sum||F_{obs}| - |F_{calc}||/\sum|F_{obs}|$, where all reflections belong to a test set of 10% data randomly selected in Phenix.

**Table S2. Cryogenic electron microscopy data collection and model statistics**

| | FhaB-CT bound to microtubules *ab initio* | FhaB-CT bound to 14-3 microtubules | FhaB-CT bound to 14-3 microtubules | FhaB-CT bound to 14-3 microtubules aligned | FhaB-CT bound to microtubule seam |
|---|---|---|---|---|---|
| accession # | EMD-49766 | EMD-49767 | PDB:9NNL EMD-49577 | EMD-49768 | EMD-49769 |
| **Data collection** | | | | | |
| microscope | Titan Krios | | | | |
| Voltage (kV) | 300.0 | | | | |
| detector | Thermo Scientific Falcon 4i | | | | |
| pixel size (Å) | 1.2 | | | | |
| total electron dose (e–/Å$^2$) | 50.0 | | | | |
| defocus range (μm) | –0.8 to –2.0 | | | | |
| energy filter slit width (eV) | 6.0 | | | | |
| # of movies | 20436.0 | | | | |
| **Reconstruction** | | | | | |
| software | cryoSPARC | | | | |
| symmetry | helical refinement (rise 81.984 Å, twist -0.609°) | helical refinement (rise 82 Å, twist -0.531°) | | | |
| selected movies | 19222.0 | | | | |
| final # of particles | 940421.0 | 1193804.0 | | 169390.0 | |
| Overall resolution (Å) | 3.2 | 2.9 | 2.4 | 3.2 | 2.9 |
| FSC threshold | 0.1 | | | | |
| map sharpening B factor (Å$^2$) | –48.28 | –56.04 | –62.69 | –28.00 | –48.11 |
| **Refinement** | | | | | |
| model composition | | | | | |
| non-hydrogen atoms | | | 22747.0 | | |
| protein residues | | | 2808.0 | | |
| ligands | | | 8.0 | | |

| | | | |
|---|---|---|---|
| root mean square deviation | | | |
| bond lengths (Å) | | | 0.0 |
| bond angles (°) | | | 1.1 |
| validation | | | |
| MolProbity score | | | 1.2 |
| clashscore | | | 3.5 |
| poor rotamers (%) | | | 1.0 |
| Ramachandran plot (%) | | | |
| favored | | | 97.6 |
| allowed | | | 2.4 |
| outlier | | | 0.0 |

**Table S3. Polar interactions (< 4.0 Å) between FhaB-CT and microtubules**

| FhaB residue | tubulin chain | tubulin residue |
|---|---|---|
| Arg3616-MC | α (A) | Asp431-SC |
| Arg3616-SC | α (A) | Asp424-SC |
| His3617-SC | α (A) | Asp431-SC |
| His3617-SC | α (A) | Glu434-SC |
| Gln3620-SC | α (A) | Arg264-SC |
| Gln3627-SC | α (A) | Glu196-MC |
| Ser3631-SC | α (A) | Gly162-MC |
| Asp3632-SC | α (A) | Lys163-SC |
| Asn3635-SC | α (A) | Asp160-MC |
| Arg3703-SC | α (A) | Glu155-SC |
| Arg3703-SC | α (A) | His197-SC |
| Tyr3706-SC | α (A) | Glu196-SC |
| Tyr3706-SC | α (A) | Arg264-SC |
| Tyr3706-SC | α (A) | Asp424-SC |
| Thr3708-SC | α (A) | Glu420-SC |
| Thr3708-SC | α (A) | Glu423-SC |
| Lys3710-SC | α (A) | Glu423-SC |
| Arg3628-SC | β (B) | Gly400-MC |
| Thr3683-SC | β (B) | Arg391-MC/SC |
| Lys3637-SC | α (E) | Lys338-MC |
| Asn3701-SC | α (E) | Arg339-SC |

Abbreviations: MC, main chain; SC, side chain

**Table S4. Bacterial strains.**

| ***strain*** | ***description*** | ***reference*** |
|---|---|---|
| *B. bronchiseptica* RB50 | wild-type *Bordetella bronchiseptica* BAA-588 | ATCC |
| *B. pertussis* Tohama I | wild-type *Bordetella pertussis* Tohama I (CIP 81.32) | (*65)* |
| CH1164 | RB50 Δ*fhaB* | This study |
| CH1174 | RB50 *fhaB(Q2721stop)* | This study |
| CH1178 | RB50 *fhaB(ΔP3386-P3590)* | This study |
| CH1195 | RB50 Δ*fimABCD* | This study |
| CH1198 | RB50 *fhaB(ΔV2016-T2274)* | This study |
| CH1222 | RB50 *fhaB(ΔL2469-T2595)* | This study |
| CH1223 | RB50 Δ*cyaA fhaB(ΔL2469-T2595)-CT*$^{Ykris}$/*cdiI*$^{Ykris}$ | This study |
| CH1224 | RB50 *fhaB(ΔH1147-G1377)* | This study |
| CH1228 | RB50 *fhaB(ΔA567-A680)* | This study |
| CH1238 | RB50 *fhaB(R1211G,D1213G)* | This study |
| CH1271 | RB50 Δ*fhaB-fimABCD-fhaC* | This study |
| CH1329 | RB50 Δ*fhaC* | This study |
| CH1338 | RB50 *fhaB(ΔT2766-L2863)* | This study |
| CH1344 | RB50 Δ*cyaA fhaB(ΔT2766-L2863)-CT*$^{Ykris}$/*cdiI*$^{Ykris}$ | This study |
| CH1346 | RB50 Δ*cyaA fhaB-CT*$^{Ykris}$/*cdiI*$^{Ykris}$ Δ*fhaC* | This study |
| CH1421 | RB50 Δ*cyaA* | This study |
| CH1422 | RB50 Δ*cyaA fhaB-CT*$^{Ykris}$/*cdiI*$^{Ykris}$ | This study |
| CH1546 | RB50 Δ*cyaA fhaB-CT(H175A)*$^{Ykris}$/*cdiI*$^{Ykris}$ | This study |
| CH1547 | RB50 Δ*cyaA fhaB(ΔV2016-T2274)-CT*$^{Ykris}$/*cdiI*$^{Ykris}$ | This study |
| CH1550 | RB50 Δ*cyaA fhaB-CT*$^{Ykris}$/*cdiI*$^{Ykris}$ Δ*fimABCD* | This study |
| CH1551 | RB50 Δ*cyaA fhaB(ΔA567-A680)-CT*$^{Ykris}$/*cdiI*$^{Ykris}$ | This study |
| CH1573 | RB50 *fhaB(ΔT3610-K3710)* | This study |
| CH1639 | RB50 *fhaB(ΔG464-G975)* | This study |
| CH1640 | RB50 Δ*cyaA fhaB(ΔG464-G975)-CT*$^{Ykris}$/*cdiI*$^{Ykris}$ | This study |
| CH1668 | RB50 *fhaB(ΔA1918-Q1969)* | This study |
| CH1669 | RB50 Δ*cyaA fhaB(ΔA1918-Q1969)-CT*$^{Ykris}$/*cdiI*$^{Ykris}$ | This study |
| CH1670 | RB50 Δ*cyaA fhaB(ΔA1918-Q1969/ΔL2469-T2595)-CT*$^{Ykris}$/*cdiI*$^{Ykris}$ | This study |
| CH1699 | RB50 *fhaB(ΔA1918-T2274)* | This study |
| CH1700 | RB50 Δ*cyaA fhaB(ΔA1918-T2274)-CT*$^{Ykris}$/*cdiI*$^{Ykris}$ | This study |
| CH2016 | *E. coli* X90 (DE3) Δ*rna* Δ*slyD::kan*, Kan$^r$ | (*46)* |
| CH2133 | RB50 Δ*cyaA fhaB(R1211G,D1213G)-CT*$^{Ykris}$/*cdiI*$^{Ykris}$ | This study |

| | | |
|---|---|---|
| CH2298 | RB50 Δ*cyaA fhaB(ΔP3386-P3590)-CT*$^{Ykris}$*/cdiI*$^{Ykris}$ | This study |
| MFD*pir*+ | *E. coli* MG1655 *RP4-2-Tc::[ΔMu1::aac(3)IV-ΔaphA-Δnic35-ΔMu2::zeo] ΔdapA::(erm-pir) ΔrecA*, Ap$^r$ Zeo$^r$ | (*43*) |
| JH_1101 | CIP 81.32 Δ*fhaB* | (*9*) |
| JH_1102 | CIP 81.32 *fhaB(ΔG2880-K3590)* | This study |
| JH_1103 | CIP 81.32 *fhaB(ΔL1871-A2358)* | This study |
| JH_1105 | CIP 81.32 *fhaB(ΔT2359-T2485)* | This study |
| JH_1106 | CIP 81.32 *fhaB(ΔT2656-L2753)* | This study |
| JH_1107 | CIP 81.32 *fhaB(ΔP3276-K3590)* | This study |
| JH_1108 | CIP 81.32 *fhaB(ΔL3493-K3590)* | This study |
| JH_1101 | CIP 81.32 *fhaB*$^+$ | This study |
| Abbreviations: Ap$^r$, apramycin resistant; Kan$^r$, kanamycin resistant; Str$^r$, streptomycin resistant; Zeo$^r$, zeocin resistant; | | |

**Table S5. Plasmids.**

| ***plasmid*** | ***description*** | ***reference*** |
|---|---|---|
| pCH752 | pSS4245-Δ*fhaB*, Amp$^r$ Kan$^r$ | this study |
| pCH763 | pSS4245-*fhaB-CT*$^{Ykris}$*/cdiI*$^{Ykris}$, Amp$^r$ Kan$^r$ | this study |
| pCH1165 | pSH21-*fhaB(P3592-K3710)*$^{RB50}$, Amp$^r$ | this study |
| pCH1166 | pSH21-*fhaB(P3610-K3710)*$^{RB50}$, Amp$^r$ | this study |
| pCH1170 | pSS4245-*fhaB(Q2721stop)*, Amp$^r$ Kan$^r$ | this study |
| pCH1252 | pBBR1::*tdTomato*, Tp$^r$ | this study |
| pCH1318 | pSS4245-*fhaB(ΔT2766-L2863)*, Amp$^r$ Kan$^r$ | this study |
| pCH1319 | pSS4245-*fhaB(ΔH1147-G1377)*, Amp$^r$ Kan$^r$ | this study |
| pCH1320 | pSS4245-*fhaB(ΔA567-A680)*, Amp$^r$ Kan$^r$ | this study |
| pCH1326 | pSS4245-Δ*fhaC,* Amp$^r$ Kan$^r$ | this study |
| pCH1330 | pSS4245-Δ*fhaB-fimABCD-fhaC,* Amp$^r$ Kan$^r$ | this study |
| pCH1352 | pSS4245-*fhaB(ΔP3386-P3590)*, Amp$^r$ Kan$^r$ | this study |
| pCH1353 | pSS4245-Δ*fimABCD*, Amp$^r$ Kan$^r$ | this study |
| pCH1354 | pSS4245-*fhaB(ΔV2016-T2274)*, Amp$^r$ Kan$^r$ | this study |
| pCH1355 | pSS4245-*fhaB(ΔL2469-T2595)*, Amp$^r$ Kan$^r$ | this study |
| pCH1509 | pBBR1::*mNeon*, Tp$^r$ | this study |
| pCH1536 | pSS4245-Δ*cyaA,* Amp$^r$ Kan$^r$ | this study |
| pCH1543 | pSS4245-*fhaB(ΔA1918-T2274)*, Amp$^r$ Kan$^r$ | this study |
| pCH1565 | pSS4245-*fhaB(ΔT3610-K3710)*, Amp$^r$ Kan$^r$ | this study |
| pCH1637 | pSS4245-*fhaB(ΔG464-G975)*, Amp$^r$ Kan$^r$ | this study |
| pCH1638 | pSS4245-*fhaB(ΔA1918-Q1969)*, Amp$^r$ Kan$^r$ | this study |
| pCH1784 | pSH21-*fhaB(A2461-S2752)*$^{SST3}$, Amp$^r$ | this study |
| pCH1785 | pSH21-*fhaB(G2409-R2674)*$^{Yfred}$, Amp$^r$ | this study |
| pCH1811 | pSH21-*fhaB(A3374-K3710)*$^{RB50}$, Amp$^r$ | this study |
| pCH1847 | pET21-*fhaB(G73-E440)*$^{RB50}$, Amp$^r$ | this study |
| pCH2287 | pSS4245-*fhaB(ΔP3386-P3590)-CT*$^{Ykris}$*/cdiI*$^{Ykris}$, Amp$^r$ Kan$^r$ | this study |
| pCH2475 | pSS4245-*fhaB(R1211G,D1213G)*, Amp$^r$ Kan$^r$ | this study |
| pCW57.1 | Gateway destination plasmid for mammalian expression, Amp$^r$ | Addgene |
| pFBK001 | pENTRY derivative | |
| pINDUCER21 | Gateway destination plasmid for mammalian expression, Amp$^r$ | (*45*) |
| pMSC156 | pCW57.1-FLAG-his6-eGFP-*fhaB(V3553-K3710)*$^{RB50}$, Amp$^r$ | this study |
| pMSC158 | pCW57.1-FLAG-his6-eGFP-*fhaB(A3602-K3710)*$^{RB50}$, Amp$^r$ | this study |
| pMSC218 | pFBK001-*cdiI*$^{Ykris}$-FLAG | this study |
| pMSC224 | pINDUCER21-*cdiI*$^{Ykris}$-FLAG | this study |
| pMSC263 | pCW57.1-FLAG-his6-eGFP-*fhaB(A3374-K3710)*$^{RB50}$, Amp$^r$ | this study |

| | | |
|---|---|---|
| pSS4245 | mobilizable allelic exchange plasmid that expresses I-SceI endonuclease from *ptxA* promoter, $Amp^r$ $Kan^r$ | (*44*) |
| Abbreviations: $Amp^r$, ampicillin resistant; $Kan^r$, kanamycin resistant; $Tp^r$, trimethoprim resistant | | |

**Table S6. Oligonucleotides.**

| ***identifier*** | ***description*** | ***sequence*** |
|---|---|---|
| CH824 | trc-rev-seq | 5´ - CGT TCT GAT TTA ATC TGT ATC AGG C |
| CH5498 | RB50-fhaB-G73-Nco-for | 5´ - ACG CCA TGG GCT TGG TTC CTC AGG GG |
| CH5499 | RB50-fhaB-E440-Xho-rev | 5´ - CAC CTC GAG CCG TCC C |
| CH5502 | fhaB-A3374-Spe-for | 5´ - TTC ACT AGT GCA GGC AAG TCA CCG AAG AAG |
| CH5503 | fhaB-Xho-rev | 5´ - TTT CTC GAG CTA TTT GTT GGT TTC ATA GAA AAC CCG G |
| CH5605 | pTrc-Nsi-for | 5´ - TTT ATG CAT GAG CTC ACT AGT GGA TCC TGT TTT TTG CGC CG |
| CH5622 | SST3-fhaB-A2461-Spe-for | 5´ - TAC ACT AGT GCC GAT CCG GGG ATT CG |
| CH5623 | SST3-fhaB-Xho-rev | 5´ - CCT CTC GAG TTA GCT GTA CTG CTC GTC GGT G |
| CH5624 | Yfred-fhaB-G2409-Spe-for | 5´ - TTT ACT AGT GGC GGT TTA TCG AGT TCA AAT AA |
| CH5625 | Yfred-fhaB-Xho-rev | 5´ - CTA CTC GAG GCT ATC GTT TGG ACG AAT CC |
| CH5669 | RB50-fhaB-T3610-Spe-for | 5´ - AAG ACT AGT CCC TTG AGC GGG CGC |
| CH5671 | fhaB-P3592-Spe-for | 5´ - TTT ACT AGT CCC GCA CCG AAG CCC AAG CCC |
| CH5676 | fhaB-A3149-Xba-for | 5´ - GGA TCT AGA TGG CCG CAG CGA ACG |
| CH5677 | fhaB-G3615-Kpn-rev | 5´ - GTG GGT ACC GCT CAA GGG CGT CGT C |
| CH5681 | Ykris-G160-Kpn-for | 5´ - CCT GGT ACC GGA TTA GCT GCT CAT G |
| CH5682 | Ykris-cdiI-Sac-rev | 5´ - TTT GAG CTC AGT TCA TTT GTT TAT ATA TAT GTT TCA ACC |
| CH5717 | fhaB-A2591-Kpn-for | 5´ - CCC GGT ACC GCG CCC GCA CCG AAG CCC AA |
| CH5718 | fhaB-Sac-rev | 5´ - TTT GAG CTC TAT TTG TTG GTT TCA TAG AAA ACC CGG TAG |
| CH5720 | fhaB-R3385-Kpn-rev | 5´ - TTT GGT ACC GCG GAC CTG GTT CTT CTT CTT CGG |
| CH5775 | fhaB-I2223-Xba-for | 5´ - CGA TCT AGA CTG GCC GGG TCA TGC TGG |
| CH5776 | fhaB-Q2721stop-Sac-rev | 5´ - ATA GAG CTC TAG CCG CCT TGG GCA TCC AGG G |
| CH5789 | fhaC-Sac-for | 5´ - TTT GAG CTC CAA CAT GAC TGA CGC AAC GAA CCG TTT C |

| | | |
|---|---|---|
| CH5790 | fhaC-D312-Bam-rev | 5´ - TGA GGA TCC AGG CGA GTA CGC CC |
| CH5807 | ΔfhaC-Spe-for | 5´ - TAG ACT AGT ATT TCT GGT CGA GCA CCT CGC |
| CH5808 | ΔfhaC-Sac-rev | 5´ - AAA GAG CTC GTT GCG TCA GTC ATA GTT CAA GG |
| CH5809 | ΔfhaC-Sac-for | 5´ - TCT GAG CTC GGC GAG AAC GCG GC |
| CH5810 | ΔfhaC-Bam-rev | 5´ - GCC GGA TCC TGC ACA CCA ACG TCA TC |
| CH5811 | fhaB-S1684-Spe-for | 5´ - ACG ACT AGT GGG CAG GCG GAC AAT CGG |
| CH5812 | fhaB-R2015-Sac-rev | 5´ - ACG GAG CTC CGA TTG TCC ACG TCG CCG CCG |
| CH5813 | fhaB-V2275-Sac-for | 5´ - TCG GAG CTC CGT GCT GGC CGC CGG CG |
| CH5814 | fhaB-T2603-Eco-rev | 5´ - TAC GAA TTC GTG ATG CCC TGG CGC GTC G |
| CH5817 | fhaB-L2158-Spe-for | 5´ - ATG ACT AGT TGG GCC AAC GTT ATG GTA AGG |
| CH5820 | fhaB-D2920-Bam-rev | 5´ - CCG GGA TCC AGG ACA TTG CCA GCC TCG |
| CH5855 | fhaB-A2468-Eco-rev | 5´ - CGC GAA TTC GCG ACG GCT GGC GGC G |
| CH5856 | fhaB-E2596-Eco-for | 5´ - GTA GAA TTC CGA GGC GAC GCG CCA GGG |
| CH5857 | fhaB-G204-Not-for | 5´ - ACG GCG GCC GCA TCG GCC TTG AT |
| CH5858 | fhaB-A566-Eco-rev | 5´ - CCG GAA TTC GCC GAA CCC AGC GTG ACG G |
| CH5859 | fhaB-N681-Eco-for | 5´ - CGC GAA TTC CGA TGC GGA ACT GCA CGT GTC |
| CH5860 | fhaB-S977-Bam-rev | 5´ - CGT GGA TCC GCC GCG CAC GTC GA |
| CH5861 | fhaB-G817-Spe-for | 5´ - GAA ACT AGT CGG CGC CAT GAC CGT GAA CG |
| CH5862 | fhaB-G1131-Eco-rev | 5´ - ACG GAA TTC CCC GAG AGC TCC AGC GCC |
| CH5863 | fhaB-V1379-Eco-for | 5´ - GAC GAA TTC CGT GAT GAA CAA GGG CTA CAT CTC G |
| CH5864 | fhaB-T1700-Sbf-rev | 5´ - GGG CCT GCA GGG TGA AAT CGT GGC CGG CC |
| CH5928 | fhaB-GGG-rev | 5´ - ACG CCC TGA TGC GGA CCG CCG CCC CCG ACC GTG ACA T |
| CH5929 | fhaB-GGG-for | 5´ - ATG TCA CGG TCG GGG GCG GCG GTC CGC ATC AGG GCG T |
| CH5936 | mNeon-Nco-for | 5´ - CCA CCA TGG TGA GCA AGG GCG |
| CH5937 | mNeon-kill-rev | 5´ - GGA GCC ATC TAC CAT AGC GGC CTG GAA AGG C |
| CH5938 | mNeon-kill-for | 5´ - GCC TTT CCA GGC CGC TAT GGT AGA TGG CTC C |

| | | |
|---|---|---|
| CH5939 | mNeon-Xho-rev | 5´ - TTT CTC GAG TTA CTT GTA CAG CTC GTC CAT GCC C |
| CH6121 | ΔcyaA-Spe-for | 5´ - AGG ACT AGT GTT ACG GCC GCG |
| CH6122 | ΔcyaA-Sac-rev | 5´ - TTT GAG CTC ACC CTA CTC GGA CAG GAA AAT C |
| CH6123 | ΔcyaA-Sac-for | 5´ - CTG GAG CTC CGT GGA TCA CGG CCC |
| CH6124 | ΔcyaA-Bam-rev | 5´ - GTC GGA TCC AGC GAC AGA AGA AAA TAG CAA GG |
| CH6214 | fhaB-R2471-Not-for | 5´ - GCC GCG GCC GCT GTT CGA AAC CCG |
| CH6215 | fhaB-S2765-Sac-rev | 5´ - GTC GAG CTC TTG TTC TCG TAG CTG GAC CCC G |
| CH6216 | fhaB-G2864-Sac-for | 5´ - TTT GAG CTC TGG CAG CGA AAA AGG TCT GGA GGC GC |
| CH6217 | fhaB-G3145-Bam-rev | 5´ - CCG GGA TCC CGG CAT AGG TCT TGC CGG TTT CGC |
| CH6218 | fhaB-V3276-Not-for | 5´ - TCG GCG GCC GCG TCA ATG CGC AAA ATC TG |
| CH6219 | fhaB-T3610amb-Sac-rev | 5´ - AAG GAG CTC TAC TTG CCC GGC TTC GGA CGC |
| CH6221 | human 18S probe | 5´ - CGG AAC TAC GAC GGT ATC TG |
| CH6244 | fhaB-G463-Bam-rev | 5´ - GGC GGA TCC GGC ACG CAC CGA CA |
| CH6245 | fhaB-S977-Bam-for | 5´ - GGC GGA TCC ACG GTG GCG GCG AA |
| CH6246 | fhaB-P1305-Avr-rev | 5´ - GCG CCT AGG CAT GGT TTC CTC GTT |
| CH6259 | fhaB-V1577-Spe-for | 5´ - TCT ACT AGT GCG CGA GGG CGT GCG C |
| CH6260 | fhaB-T1917-Sac-rev | 5´ - AGC GAG CTC GTG TTC TCG ATG CGC GGC GC |
| CH6261 | fhaB-S1971-Sac-for | 5´ - CAC GAG CTC GCT CAT CGA GGT CGG CAA G |
| CH6262 | fhaB-K2306-Eco-rev | 5´ - CCG GAA TTC AGG TTT TCC GGC CGG CCG |
| CH6303 | human 18S 5´-probe | 5´ - GCT ACT GGC AGG ATC AAC CAG GTA |
| CH6305 | human 18S 3´-probe | 5´ - CGG AAA CCT TGT TAC GAC TTT TAC TTC C |
| CH6325 | PAPK-Ykris-rev | 5´ - CCG CTT CAT GAG CAG CTA ATC CCC CGC TCA AGG GCG TCG TCT |
| CH6328 | Ykris-G160-for | 5´ - GGA TTA GCT GCT CAT GAA GCG G |
| MSC33 | pCW57-fhaB-rev | 5´ - TCT AGA GTC GCG GCC GCC TAT TTG TTG GTT TCA TAG AAA ACC CGG T |
| MSC37 | pCW57-FLAG-for | 5´ - AAG CAG GCT CGC TAC CGG TCG CCA CCA TGG ACT ACA AGG ACC ATG ACG GTG A |
| MSC38 | GFP-rev | 5´ - TTT GTA TAG TTC ATC CAT GCC ATG TGT |

| | | |
|---|---|---|
| MSC39 | GFP-fhaB(V3553)-for | 5´ - GAT GAA CTA TAC AAA ggc ggg tcg GTG GCG GAG GCT GGC A |
| MSC40 | GFP-fhaB(A3602)-for | 5´ - GAT GAA CTA TAC AAA GGC GGG TCG GCC GAG CGT CCG AAG C |
| MSC66 | ΔfhaB-Not-for | 5´ - GGG GCG GCC GCT CAG CTT CTG CAG CAG GCG |
| MSC67 | ΔfhaB-Sac-rev | 5´ - CCC GAG CTC ATT CCG ACC AGC GAA GTG AAG T |
| MSC68 | ΔfhaB-Sac-for | 5´ - GGG GAG CTC GTA GTC GCT GCC CGC C |
| MSC69 | ΔfhaB-Bam-rev | 5´ - CCC GGA TCC TGA CAA GCC GAC CAT CCC G |
| MSC81 | diI_FL_R | 5´ - CTT GTC ATC GTC ATC CTT GTA GTC CGA TCC GTT CAT TTG TTT ATA TAT ATG TTT CAA CCA |
| MSC108 | tomato-Nco-for | 5´ - ACA CAG GAA ACA GAC CAT GGT GAG CAA GGG CGA G |
| MSC109 | tomato-Xho-rev | 5´ - ACC GAT CGG GTA CCC TCG AGT TAC TTG TAC AGC TCG TCC ATG C |
| MSC273 | Ykris-cdiI-FLAG-for | 5´ - ACA AGG ATG ACG ATG ACA AGT AAG CGG CCG CAT CGA TTC T |
| MSC299 | GFP-fhaB(A3374)-for | 5´ - GAT GAA CTA TAC AAA GGC GGG TCG GCA GGC AAG TCA CCG AAG AA |
| | Bp-ΔfhaB-Not-for | 5´ - CTG CGG CCG CGG CAT TGA TGA CCT CGT GCA G |
| | Bp-ΔfhaB-Spe-rev | 5´ - GAA CTA GTC GTG TTC ATA TTC CGA CCA GC |
| | Bp-ΔfhaB-Spe-for | 5´ - CTA CTA GTA ACA AAT AGG TAG TCG CGG CCT G |
| | Bp-ΔfhaB-Bam-rev | 5´ - AGG ATC CCA TGC CGC CTT GCC GCT TTA C |
| | Bp-Δex2-up-for | 5´ - CTA TGC TAG GGC GGC CGC ACA TCA ACA GCG CCA AGC TGG |
| | Bp-Δex2-up-rev | 5´ - GGC GCG ACC CGG ACC GTA TAG AGA TCC TTG CCG ACC TC |
| | Bp-Δex2-down-for | 5´ - GAG GTC GGC AAG GAT CTC TAT ACG GTC CGG GTC GCG CC |
| | Bp-Δex2-down-rev | 5´ - GAC GCG TGG ATC CGA ATT CTG AGA CGT CGA CCT CGT C |
| | Bp-Δprodomain-up-for | 5´ - CCT ATG CTA GGG CGG CCG CAG CCC GCG ACG CGA GGT T |
| | Bp-Δprodomain-up-rev | 5´ - CAG GCG CGT GGC GGC CGT CA |
| | Bp-Δprodomain-down-for | 5´ - TGA CGG CCG CCA CGC GCC TGT AGG TAG TCG CGG CCT GC |
| | Bp-Δprodomain-down-rev | 5´ - AGG ACG CGT GGA TCC GAA TTG GAG AAC CCC ACC TGA CTG |
| | Bp-ΔSA-up-for | 5´ - CTA TGC TAG GGC GGC CGC ACA AGT CGC AGA TCG ACG CGG |

| | | |
|---|---|---|
| | Bp-ΔSA-up-rev | 5´ - GAT GCC CTG GCG CGT CGC CTC CGC GAC GGC TGG CGG CGC GA |
| | Bp-ΔSA-down-for | 5´ - TCG CGC CGC CAG CCG TCG CGG AGG CGA CGC GCC AGG GCA TC |
| | Bp-ΔSA-down-rev | 5´ - GAC GCG TGG ATC CGA ATT CAT TGC CAG CCT CGA CGG AG |
| | Bp-ΔFHA2-up-for | 5´ - CTA TGC TAG GGC GGC CGC AGT TTA TCG ACC AGA GCA AAT TC |
| | Bp-ΔFHA2-up-rev | 5´ - CTT CCA GAC CGC TTT CGC TGC CCG AGC TCT TGT TCT CGT AGC TG |
| | Bp-ΔFHA2-down-for | 5´ - CAG CTA CGA GAA CAA GAG CTC GGG CAG CGA AAG CGG TCT GGA AG |
| | Bp-ΔFHA2-down-rev | 5´ - GAC GCG TGG ATC CGA ATT GAT TCG AAG GTC GAT TCG GCC |
| | Bp-ΔPRR-up-for | 5´ - CCT ATG CTA GGG CGG CCG CAG CCG GCT TCA ATT TCA ATA CG |
| | Bp-ΔPRR-up-rev | 5´ - GCG GAC CTG GTT CTT CTT CT |
| | Bp-ΔPRR-down-for | 5´ - AGA AGA AGA ACC AGG TCC GCT AGG TAG TCG CGG CCT GC |
| | Bp-ΔCT-up-for | 5´ - CCT ATG CTA GGG CGG CCG CAA AGC ACT GGG CCG GAG GC |
| | Bp-ΔCT-up-rev | 5´ - CGG CAG GCC GCG ACT ACC TAG GGC GTC GTT TTG CCC GGC |
| | Bp-ΔCT-down-for | 5´ - GCC GGG CAA AAC GAC GCC CTA GGT AGT CGC GGC CTG CCG |
| | Bp-ΔCT-down-rev | 5´ - AGG ACG CGT GGA TCC GAA TTG GAG AAC CCC ACC TGA CTG |

**Movie S1. Confocal microscopy of U2OS cells expressing GFP fused to FhaB residues Ala3374-Lys3710.** GFP-PRR-CT was induced with doxycycline for 24 h, then stained with antibodies to β-tubulin and Hoechst for visualization by confocal fluorescence microscopy. Scale bar = 10 μm.

**Movie S2. Confocal microscopy of U2OS cells expressing GFP fused to FhaB residues Val3553-Lys3710.** GFP-CT was induced with doxycycline for 24 h, then stained with antibodies to β-tubulin and Hoechst for visualization by confocal fluorescence microscopy. Scale bar = 10 μm.

**Movie S3. Confocal microscopy of U2OS cells expressing GFP fused to FhaB residues Ala3374-Lys3710.** GFP-CT was induced with doxycycline for 24 h, then stained with antibodies to β-tubulin and Hoechst for visualization by confocal fluorescence microscopy. Scale bar = 10 μm.

**Movie S4. Rotation of the consensus 14-3 microtubule map (EMD-49767).**

**Movie S5. Rotation of the aligned 14-3 microtubule map (EMD-49768).**

**Movie S6. Time-lapse microscopy of *fhaB*$^{+}$ bacteria bound to tracheal respiratory epithelia.** Scale bar = 10 μm.

**Movie S7. Time-lapse microscopy of *fhaB-∆CT* bacteria bound to tracheal respiratory epithelia.** Scale bar = 10 μm.

**Movie S8. Time-lapse microscopy of *fhaB-∆CT* bacterium shed from motile cilia.** tdTomato expressing RB50 *fhaB-∆CT* bacteria (red) were visualized on fluorescently labeled cilia. Scale bar = 10 μm.

**Movie S9. Time-lapse microscopy of *fhaB*$^{+}$ bacteria bound to tracheal respiratory epithelia.** tdTomato expressing RB50 *fhaB*$^{+}$ bacteria (red) were visualized on fluorescently labeled cilia. Scale bar = 10 μm.
